# G-quadruplex Structures in Dysregulated Long Non-Coding RNA of Ovarian Cancer and their Binding Interactions with Human Serum Albumin

**DOI:** 10.1101/2024.11.20.624450

**Authors:** Deepshikha Singh, Chinmayee Shukla, Bhaskar Datta

## Abstract

Ovarian carcinoma (OC) is a major cause of cancer-related mortality among women worldwide, with especially poor outcomes in resource-constrained regions. Long non-coding RNAs (lncRNAs) have emerged as critical players in OC progression, influencing drug resistance, metastasis, and other cellular processes. This study examines the potential of dysregulated lncRNAs in OC to form G-quadruplex (G4) structures. Using in silico predictions and experimental validation, we identified five lncRNAs ERLNC1, DLX6-AS1, LINC01127, FMNL1-DT, and LINP1 as candidates with a high propensity towards G4 formation. A combination of circular dichroism (CD) spectroscopy, Thioflavin T (ThT) fluorescence assay, Dot Blot assay, and reverse transcriptase stop assay, confirmed the ability of the lncRNAs to fold into stable G4 structures. Competitive DNA binding assays enabled the identification of essential G-tracts within each G4 motif highlighting the subtle differences in structure and stability of the different quadruplexes. Human serum albumin (HSA), a major circulatory protein, was found to interact with these G4-forming lncRNAs, albeit with different affinities and structural implications on the G4 motifs. This study provides the first detailed characterization of G4 structures in OC-dysregulated lncRNAs and elucidates their interactions with HSA. The G4 structures identified in these lncRNAs and their interaction with HSA can be of potential value for early OC diagnosis.

**Graphical Abstract:** 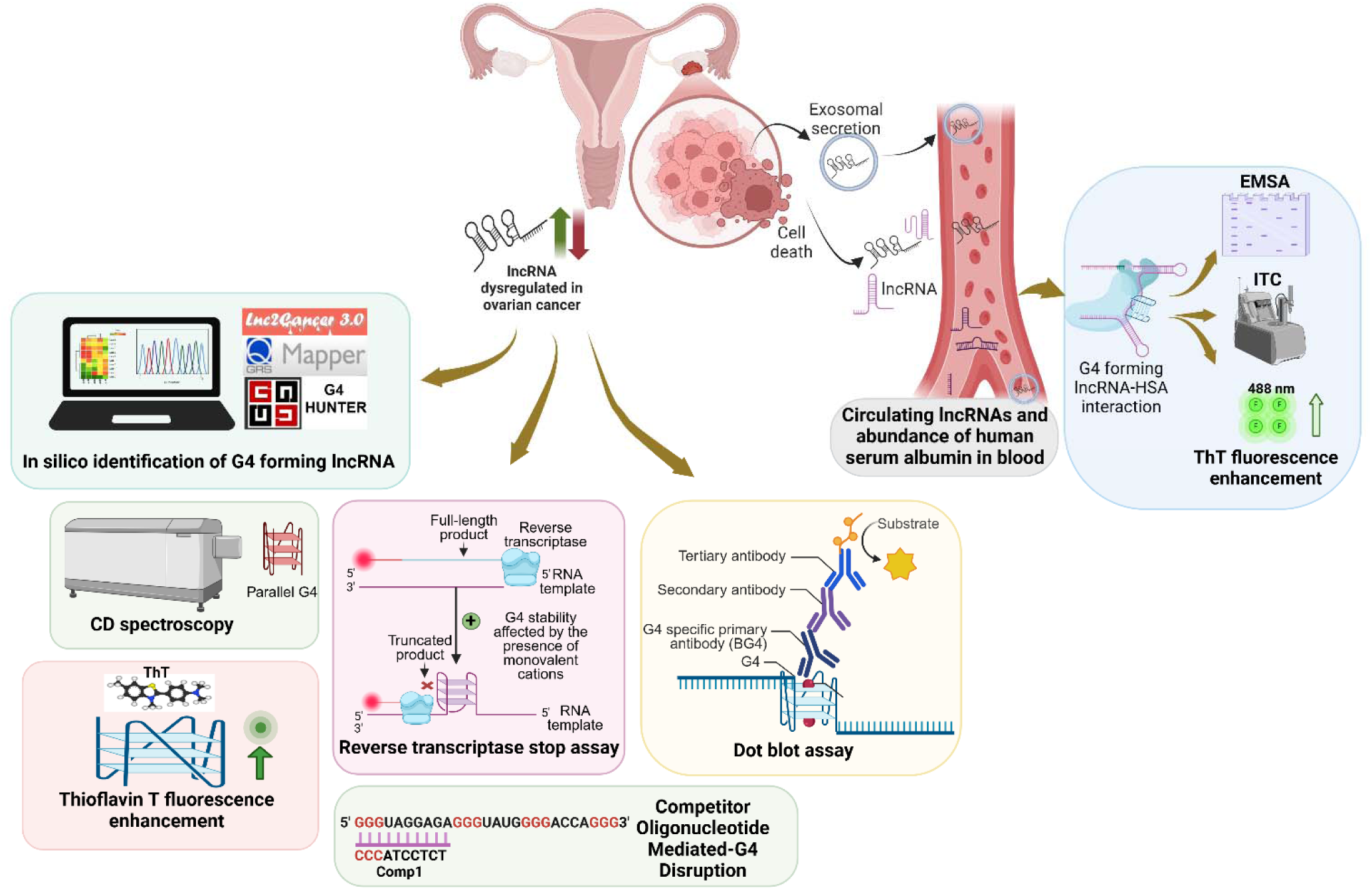

## 1. Introduction

Non-coding RNAs (ncRNAs) are recognized as key components of transcriptional and post-transcriptional control of gene expression(*1,2*). Long non-coding RNAs (lncRNAs) are longer than 500 nucleotides in length. While these were previously considered transcriptional noise, they are now recognized as key players in several cellular processes that are especially relevant to cancer progression. The dysregulation of lncRNA hampers cellular homeostasis and is associated with cancer treatment resistance, invasion, metastasis, migration, proliferation, apoptosis, altered cell metabolism and cell cycle regulation(*3–6*). LncRNA expression is more tissue- and stage-specific than protein-coding genes, suggesting that it is maintained selectively and can thereby serve as an effective biomarker and therapeutic target(*7*). The complex structures formed by lncRNAs enable access to a plethora of interacting partners, including miRNA, mRNA and proteins, ultimately resulting in substantive functional outputs. Unlike protein-coding genes, many lncRNAs show low sequence conservation across species, making functional interpretation difficult. However, their secondary structures may be conserved, suggesting that structural preservation, rather than sequence similarity, underpins their functional roles(*8,9*). The distinctive tertiary structures formed by lncRNAs arise from the interplay of secondary structural motifs such as hairpins, duplexes, bulges and stem-loops. G-quadruplexes (G4s) are among the secondary structure motifs at the disposal of lncRNAs. RNA G4s (RG4s) have been suggested to possess comparable biological significance and greater thermodynamic stability versus DNA G4s(*10*).

The systematic scrutiny of G4s as potential anchors that facilitate the association of lncRNAs with specific interacting partners could shed light on the functional roles of the lncRNA. In this work, we have identified and studied the G4s formed by several lncRNAs that are dysregulated in ovarian cancer (OC). OC is a widespread female malignancy with a relatively dismal prognosis especially in regions with resource constraints(*11*). In 2023, there were about 324,603 new cases of OC diagnosed and about 206,956 new deaths globally(*12*). The functions of several lncRNAs in OC have been scrutinized at length. The lncRNA MALAT-1 is overexpressed in OC and is linked to cell growth, migration, and invasion. MALAT-1 accelerates the progression of OC via regulation of the expression of metalloproteinases, such as MMP13 upregulation and MMP19 and ADAMTS1 downregulation(*13*). The lncRNA UCA1 is overexpressed in epithelial ovarian cancer and functions as a competitive endogenous RNA that upregulates the expression of MMP14 by directly binding to miR-485-5p. (*14*). OC tissues exhibit much higher levels of lncRNA H19 expression, and knockdown of H19 in OC cell lines reduced cell proliferation and promoted apoptosis(*15*). lncRNAs HOTAIR and ANRIL induce proliferation, migration, and invasion in OC(*16,17*). G4 structures in lncRNAs may influence disease-associated cellular functions by affecting protein interactions(*18*). It has been shown that RNA-binding proteins interact with folded and unfolded RG4s to regulate various processes. An example is hnRNP H/F protein, which regulates the translation of RG4-containing mRNAs, encoding proteins that maintain genome stability and mRNAs that code for stress response factors(*19*). The lncRNA GSEC binds to the DHX36 RNA helicase via a G4 motif, preventing its G-quadruplex unwinding activity and aiding in the migration of colon cancer cells(*20*). Notwithstanding the large number of putative G4-forming sequences harboured by lncRNAs, the existing studies are grounded in a protein-centred perspective. We have previously demonstrated an *in silico* methodology for identifying G4-bearing lncRNAs dysregulated in cervical cancer(*21*).

In this work, we have selected lncRNAs dysregulated in OC that have the potential to form G4s and used a combination of spectroscopic and molecular biology experiments to characterize and validate the *in vitro* G4 formation by the selected lncRNAs. Further, we demonstrate the distinctive and hitherto unknown ability of specific lncRNAs to bind with human serum albumin (HSA). HSA is the most abundant protein in blood plasma, with a concentration of 35–50 mg/mL(*22*). Serum albumin also has prognostic significance with lower levels of HSA being correlated with poor survival outcomes in colorectal cancer patients (*23*). Although lncRNAs and HSA coexist in circulatory fluids, information on their specific interaction is unclear. This is the first report of a distinctive interaction pattern of G4 motifs in a cancer-dysregulated lncRNAs with HSA.

## 2. Materials and methods

### 2.1. Reagents

Nuclease free water (Catalog no. 96370) and urea (Catalog no. 21113) were obtained from Sisco Research Laboratories Pvt. Ltd., Mumbai, India. Human serum albumin (Catalog no. A8763), Formamide (Catalog no. 47670), bromophenol blue (Catalog no. 114391), EDTA (Catalog no. EDS), Braco19 (Catalog no. SML0560), xylene cyanol (Catalog no. X4126), Bovine Serum Albumin (Catalog no. A9418) were obtained from Sigma-Aldrich Chemicals Pvt. Ltd., Bangalore, India. Anti-DNA G-quadruplex structures Antibody, clone BG4 (Catalog no. MABE917), DYKDDDDK Tag (D6W5B) Rabbit mAb (Catalog no. 14793), and Goat anti-Rabbit IgG (H+L) Cross-Adsorbed Secondary Antibody were procured from Sigma-Aldrich Chemicals Pvt. Ltd., Bangalore, India, Cell Signaling Technology, Inc., Research Instruments Pte. Ltd., Singapore, respectively. Carboxy pyridostatin trifluoroacetate salt (cPDS) (Catalog no. HY-112680A) was procured from MedChemExpress. All reagents were used as per the manufacturer’s protocol and stored in the recommended conditions.

### 2.2. *In silico* identification of the putative quadruplex-forming sequences in lncRNAs dysregulated in ovarian cancer

The Lnc2Cancer database was utilized to determine the lncRNAs dysregulated in OC(*24*). The aliases of all the output lncRNAs were manually compiled from GeneCards. NCBI nucleotide database was used to extract the lncRNA sequences in FASTA format. The sequences in FASTA format were fed as a query sequence in the QGRS mapper tool using the following parameters: Maximum length: 45, Min. G-tract: 2, Loop size: 0-35. QGRS Mapper uses a scoring method to identify and evaluate potential G-quadruplex sequences with the motif G_x_N_y1_G_x_N_y2_G_x_N_y3_G_x_, where G indicates guanine nucleotides in G-tetrad formation, x is the number of G repeats (at least two), and y1, y2, and y3 denote loop sequence lengths(*25*). The G4 Hunter tool was also used for G4 prediction, with a window size set at 45(*26*). The G4 Hunter score threshold of ∼1.4 was used to select 3G quadruplex forming PQS, and ∼0.9 score threshold was used for 2G quadruplex forming PQS. PQS common in both QGRS mapper and G4 Hunter were selected for further studies (**Table 2**).

### 2.3. Oligonucleotides

The oligonucleotide templates used for *in vitro* transcription are shown in **Table 1**. The oligonucleotides were bought in lyophilized form from Sigma-Aldrich Chemicals Pvt. Ltd., Bangalore, India. Stocks of 100 µM were prepared with nuclease-free water and stored at −20 °C until use. The basis of design of the oligonucleotides is provided in **Suppl. Fig. 1**.

**Table 1.**
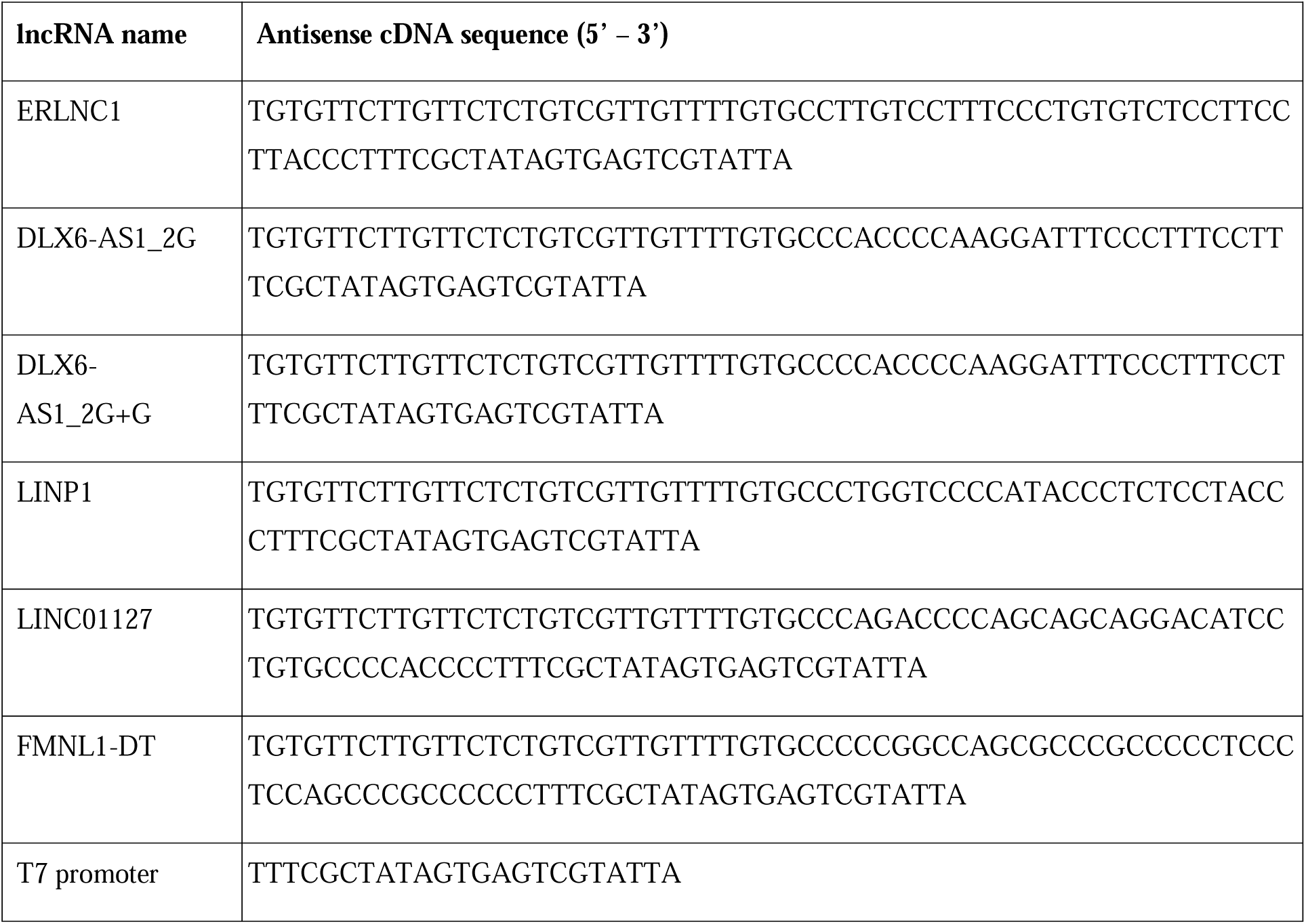
Oligonucleotide templates used for in-vitro transcription.

### 2.4. *In vitro* transcription

The sense and antisense DNA strands of the T7 RNA promoter were annealed in a buffer (termed annealing buffer) consisting of 10 mM Tris pH 8.0, 50 mM NaCl, and 1 mM EDTA pH 8.0. The concentration and purity of annealed DNA oligonucleotides were quantified using NanoDrop™ 2000 spectrophotometer (Thermo Fisher Scientific, USA). In vitro-transcription of the annealed DNA oligonucleotides was carried out using HiScribe^™^ T7 High Yield RNA Synthesis Kit (Catalog no. E2040S, New England Biolabs Pte. Ltd., Singapore), following the manufacturer’s protocol. The DNA oligonucleotides in the transcribed RNAs solution were digested using DNase I (RNase-free) (Catalog no. M0303L, New England Biolabs Pte. Ltd., Singapore). RNAs were cleaned and eluted using Monarch^®^ RNA Cleanup Kit (Catalog no. T2050L, New England Biolabs Pte. Ltd., Singapore), following the manufacturer’s protocol. The concentration and purity of eluted RNAs were quantified using NanoDrop™ 2000. RNase Inhibitor (Catalog no. M0314L, New England Biolabs Pte. Ltd., Singapore) was used to inhibit RNases and were stored at −80 °C until further use.

### 2.5. Circular Dichroism (CD) spectroscopy

Samples containing 5 μM RNA were folded in a buffer (termed folding buffer) containing 10mM Tris-Cl pH 7.5 and 0.01 mM EDTA pH 8.0, by incubating at 95 °C for 5 mins and gradually cooling to room temperature. RNA samples were supplemented with 100 mM KCl or LiCl and increasing concentrations of TMPyP4 (5, 10, 15, 20 and 25 µM). CD spectra were recorded on a JASCO J-815 spectropolarimeter using a quartz cell of 1 cm optical path length and the following parameters, Instrument scanning speed: 100 nm/min; Wavelength range: 190-320 nm; Digital Integration Time (DIT): 1 sec; Bandwidth: 1.00nm; Accumulations: 3; Temperature: 16 °C.

### 2.6. ThT Fluorescence enhancement assays

Fluorescence enhancement assays were performed using Thioflavin T (ThT) (Catalog no. T3516, Sigma-Aldrich, USA) in a 96-well black fluorescence microplate. RNA samples (2 µM) were folded in folding buffer containing 10mM Tris-Cl pH 7.5 and 0.01 mM EDTA pH 8.0 by incubating at 95 °C for 5 mins, followed by gradually cooling to room temperature. ThT (2 μM) was added to the folded RNA, and the emission spectra was obtained after excitation at 445 nm. The excitation spectra were obtained with emission captures at 488 nm. Single-point fluorescence intensities were also obtained for ThT at the mentioned wavelengths. The fluorescence of samples was measured at 25 °C using Cytation 5 Cell Imaging Multimode Reader (Agilent Technologies, USA). The mean fluorescence intensities (in arbitrary units) were extracted from the emission spectra and were plotted against the wavelength after applying the Savitzky-Golay smoothing method with a 20-point window using Origin (Pro) software. For the ThT Fluorescence enhancement assays with HSA, the RNA was first folded as described, and then incubated with the indicated concentrations of HSA for 2 hours at 37 °C. Fluorescence measurements were performed as described above. An unpaired t-test was conducted for the statistical analysis.

### 2.7. Reverse transcriptase (RT) stop assay

Texas red tagged primers were purchased from Sigma Aldrich, USA in lyophilized form, and nuclease-free water was used to prepare 100 µM solutions. Each RT-stop experiment was performed in 10 μl reaction mixtures, containing annealing buffer (10 mM Tris pH 8.0, 50 mM NaCl, and 1 mM EDTA pH 8.0), 2 μM RNA, 200 nM Texas red-tagged primer, 2 mM dNTPs (Catalog no. U1511, Promega Biotech India Pvt. Ltd, New Delhi), and increasing concentrations of KCl/LiCl (0, 50, 100, 150 mM). For the annealing of the primer, the reaction mixture was denatured by heating at 95 °C for 5 mins, followed by cooling to room temperature. 4 U/μl per reaction reverse transcriptase (Catalog no. M5301, Promega Biotech India Pvt. Ltd, New Delhi) was added with a buffer of 3 mM MgCl_2_, 10 mM Tris-Cl pH 7.5, 1 mM EDTA pH 8.0, and incubated at 37 °C for 1 hour. An equal volume of stop buffers (95% Formamide, 0.05% Bromophenol Blue, 20 mM EDTA, 0.05% Xylene cyanol) was added to stop the reaction. The products were separated on a 15% denaturing (Urea) polyacrylamide gel. The gels were imaged using ChemiDoc^TM^ MP Imaging system using the Rhodamine filter and then counterstained with Diamond^TM^ Nucleic Acid dye (Promega Corporation) to visualize the primer bands. Band intensities were quantified using ImageJ software. Ordinary one-way ANOVA was employed for statistical analysis.

### 2.8. Competitive binding of complementary DNA oligos

The experimental design of the competitive binding assay is provided in supporting information and **Suppl. Fig. 2**. This assay followed a previously established protocol in our lab(*27*). For this assay, RNA samples (2 µM) were folded in the presence of an increasing concentration of the DNA competitor in a folding buffer (10mM Tris-Cl pH 7.5 and 0.01 mM EDTA pH 8.0) by incubating at 95 °C for 5 mins, followed by gradually cooling to room temperature. The products were separated on a 15% native polyacrylamide gel, visualized on ChemiDoc™ MP Imaging system after staining with 0.5 mM ThT, and then counterstained with Diamond™ Nucleic Acid dye (Promega Corporation) to visualize competitor oligos. Band intensities were quantified using ImageJ software. Ordinary one-way ANOVA was employed for statistical analysis,

### 2.9. Dot Blot assay

The RNA samples were prepared at a concentration of 4 µM in folding buffer containing 10 mM Tris-Cl pH 7.5 and 0.01 mM EDTA pH 8.0 with or without K^+^, Li^+^, and G4-targeting ligands, ensuring the final volume was kept as low as reasonable (3 µL). The samples were then heated at 95 °C for 5 minutes, and gradually cooled to room temperature. The prepared RNAs were loaded onto a nitrocellulose membrane and dried. The RNA was crosslinked to the membrane using a UV crosslinker at 120 millijoules/cm² for two pulses. The membranes were then blocked in 1% BSA in TBST for 1 hour at RT with gentle rocking. Subsequently, the membrane was incubated with a primary antibody (BG4 containing FLAG tag) at a 1:1000 dilution in 1% BSA in TBST at 4 °C overnight with gentle rocking. The membrane was washed three times with 1X TBST for 5 minutes each. It was then incubated with rabbit anti-FLAG secondary antibody at a 1:1500 dilution in 3% BSA in TBST for 1 hour with gentle rocking. The membrane was rewashed thrice with 1X TBST for 5 minutes each with rocking. Following this, the membrane was incubated with an anti-rabbit HRP-linked tertiary antibody at a 1:3000 dilution in 1X TBST with gentle rocking. The membrane underwent a final wash with 1X TBST thrice for 5 minutes each with rocking. Visualization was performed using BioRad ECL reagents and the ChemiDoc^TM^ MP Imaging system.

### 2.10. Electrophoretic Mobility shift assay

The primer binding site was used to label the synthesized lncRNAs with Texas-red labeled primer for visualizing electrophoretic mobility shift assays. 4 µM of RNA was first folded and annealed with the labeled primer by incubating at 95 °C for 5 mins, followed by gradually cooling to room temperature. The primer-annealed RNA was then incubated with different concentrations of HSA for 2 hours at 37 °C in 10 μL of total reaction volume. HSA solutions were prepared in a folding buffer (10 mM Tris-Cl pH 7.5 and 0.01 mM EDTA pH 8.0) and 5% glycerol. Binding reactions were loaded onto 8% polyacrylamide gels in 1X Tris-Borate EDTA. The gels were imaged using ChemiDoc^TM^ MP Imaging system using the Rhodamine filter and then counterstained with 0.5 mM ThT to visualize the G4 forming bands. Band intensities were quantified using ImageJ software. Ordinary one-way ANOVA was employed for statistical analysis.

### 2.11. Isothermal Titration Calorimetry (ITC)

ITC measurements were conducted using an ITC200 microcalorimeter (Microcal-200, Malvern Panalytical) at 25 °C. All lncRNA samples were folded in the folding buffer (10 mM Tris-Cl pH 7.5 and 0.01 mM EDTA pH 8.0) prior to titrations. The revolving syringe (750 rpm) was loaded with 20 μM HSA dissolved in the same buffer, and 5 μM lncRNA solution was injected inside the cell. Twenty injections of 40 μL of HSA were made into the RNA, with a 150 s gap between each injection. To accommodate syringe diffusion into the cell during equilibration, the first injection was 0.4 μL, and data fitting did not include this injection. The isotherms were analyzed using a single-site binding model in the Microcal ITC software to determine the relevant thermodynamic parameters.

### 2.12. Statistical Analysis

The statistical analyses included appropriate tests, and the resulting statistical significance (*P*-values) with *P* ≤ 0.05, *P* ≤ 0.01, *P* ≤ 0.001, and *P* ≤ 0.0001 are denoted with one asterisk (*), two asterisks (**), three asterisks (***), and four asterisks (****), respectively. Non-significant *P*-values are not represented.

## 3. Results and Discussion

### 3.1. lncRNAs dysregulated in ovarian cancer form G-quadruplexes

We employed an in silico G4-prediction pipeline previously established by our laboratory and named as CanLncG4 platform (https://www.canlncg4.com), which provides comprehensive predictions of G4-forming potential within lncRNAs(*28*). Briefly, we used Lnc2Cancer to identify the dysregulated lncRNAs in OC. Lnc2Cancer shows 367 entries of lncRNAs related to OC. Among these, 265 lncRNAs are upregulated, 92 lncRNAs are downregulated, and 10 display differential expression in OC. We then used the QGRS mapper and G4hunter to shortlist the lncRNAs that have strong potential to form G-quadruplexes. After removing redundant entries and analyzing all the lncRNAs, 47, 61, and 1 lncRNAs contained at least one PQS with the potential to form 2G-G4, 3G-G4, and 4G-G4 structures, respectively. Here 2G, 3G and 4G correspond to the number of consecutive G’s that constitute each putative G-tract. 4 sequences were shortlisted by comparing the QGRS mapper G score and the G4 hunter score. The 4 shortlisted candidates were selected to represent 2G and 3G bearing sequences that are reported (i) solely in the context of OC, and (ii) across other cancers. The shortlisted lncRNAs are ERLNC1 (2G), DLX6-AS1 (2G), LINC01127 (3G), LINP1 (3G) and FMNL1-DT (3G). Due to the presence of a terminal G in case of DLX6-AS1 (sequence named as DLX6-AS1_2G), we prepared another sequence with an additional G appended to the 3’ end (sequence named DLX6-AS1_2G+G). We have referred to these PQSs hereafter by the name of their respective lncRNAs. **Table 2** describes the PQS corresponding to the selected lncRNAs along with the cognate QGRS and G4hunter rating scores.

**Table 2.**
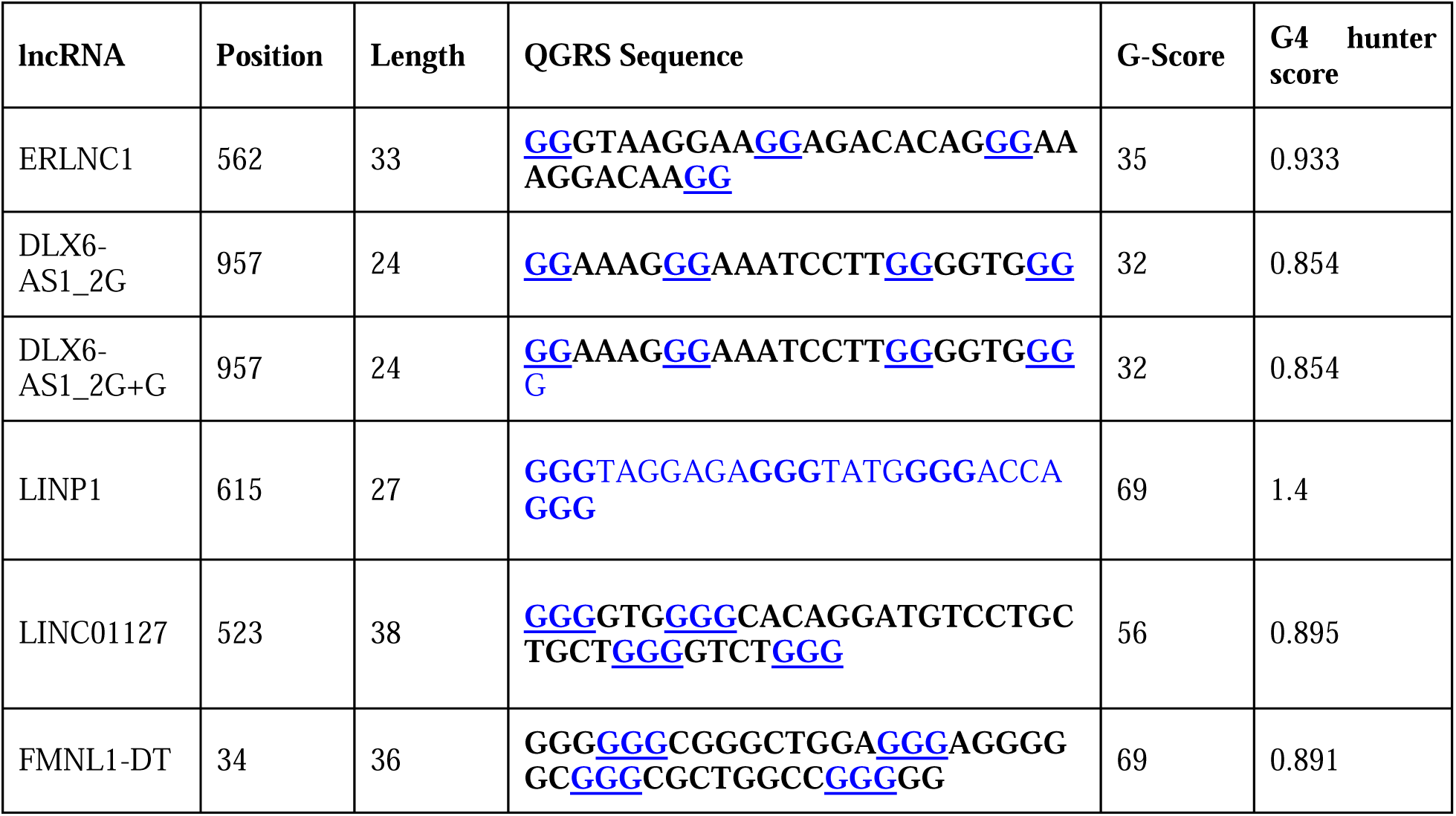
Selected lncRNAs that are dysregulated in ovarian cancer with designated PQS.

### 3.2. *In vitro* assessment of G4 formation by OC dysregulated lncRNAs

We used a combination of CD spectroscopy, ThT fluorescence enhancement assay, and Reverse Transcriptase (RT) stop assay to investigate the behavior of the selected PQS of OC dysregulated lncRNAs listed in **Table 2**. The CD spectra of ERLNC1, DLX6-AS1, DLX6-AS1_2G+G, LINP1, LIN01127 and FMNL1-DT show positive bands at ∼265 nm and negative bands at ∼245 nm. These CD signals are unambiguously indicative of parallel G-quadruplex conformations(*29,30*). We conducted CD measurements in the absence of additional monovalent ions to assess the inherent stability and conformation of the G-quadruplex structures formed by the lncRNAs. This approach allowed us to evaluate the intrinsic folding characteristics of the RNA sequences without cations’ stabilizing/destabilizing effects, providing a clearer understanding of their baseline G4-forming potential. All the lncRNAs retain the CD spectrum in the absence of additional K^+^ or Li^+^ ions. We use the term additional to indicate intentionally added ions as opposed to that which may have entered the RNA solutions as artifacts of th sample preparation protocol being followed.

The CD spectra of the RNAs display only modest changes in band intensities in the presence of K^+^ compared to the presence of Li^+^ **(Fig. 1)**. Li^+^ does not have any physiological relevance, and the purpose of using it in these experiments was solely to assess effects of a cation that i generally understood to destabilize G4s. LINP1, LINC01127, and FMNL1-DT, which form 3G-G4 structures, exhibit relatively higher CD intensities compared to DLX6-AS1, a 2G-G4. Interestingly, ERLNC1, also a 2G-G4, displays CD intensity comparable to the 3G-G4s. The 3G bearing RNAs LINC01127 followed by FMNL1-DT display sharper CD bands compared to the other sequences. Such variations in the observed CD spectra can be attributed to subtle differences in topology and stabilities of the cognate G4s(*31–34*). The behavior of PQS’ in the presence of G4-interacting ligands sheds light on similarities and differences in their conformations. We measured the CD spectra of the lncRNAs in the presence of TMPyP4, a porphyrin-derivative that is known to stabilize DNA G4’s and destabilize RNA G4s(*35–37*). A shown in **Fig. 2**, increasing concentrations of TMPyP4 resulted in variable extent of disruption in the CD spectrum of each lncRNA. Notably, significant flattening is observed in the CD spectrum of DLX6-AS1_2G and DLX6-AS1_2G+G at higher concentrations of TMPyP4. In sharp contrast, ERLNC1, LINP1, FMNL-DT1 and LINC01127 display only modest decrease in the maximum and minimum CD band intensities at 265 nm and 245 nm, respectively. The observed decrease in CD band intensities could be attributed to disruption or unfolding of th RNA G4(*38*).

**Fig. 1.**
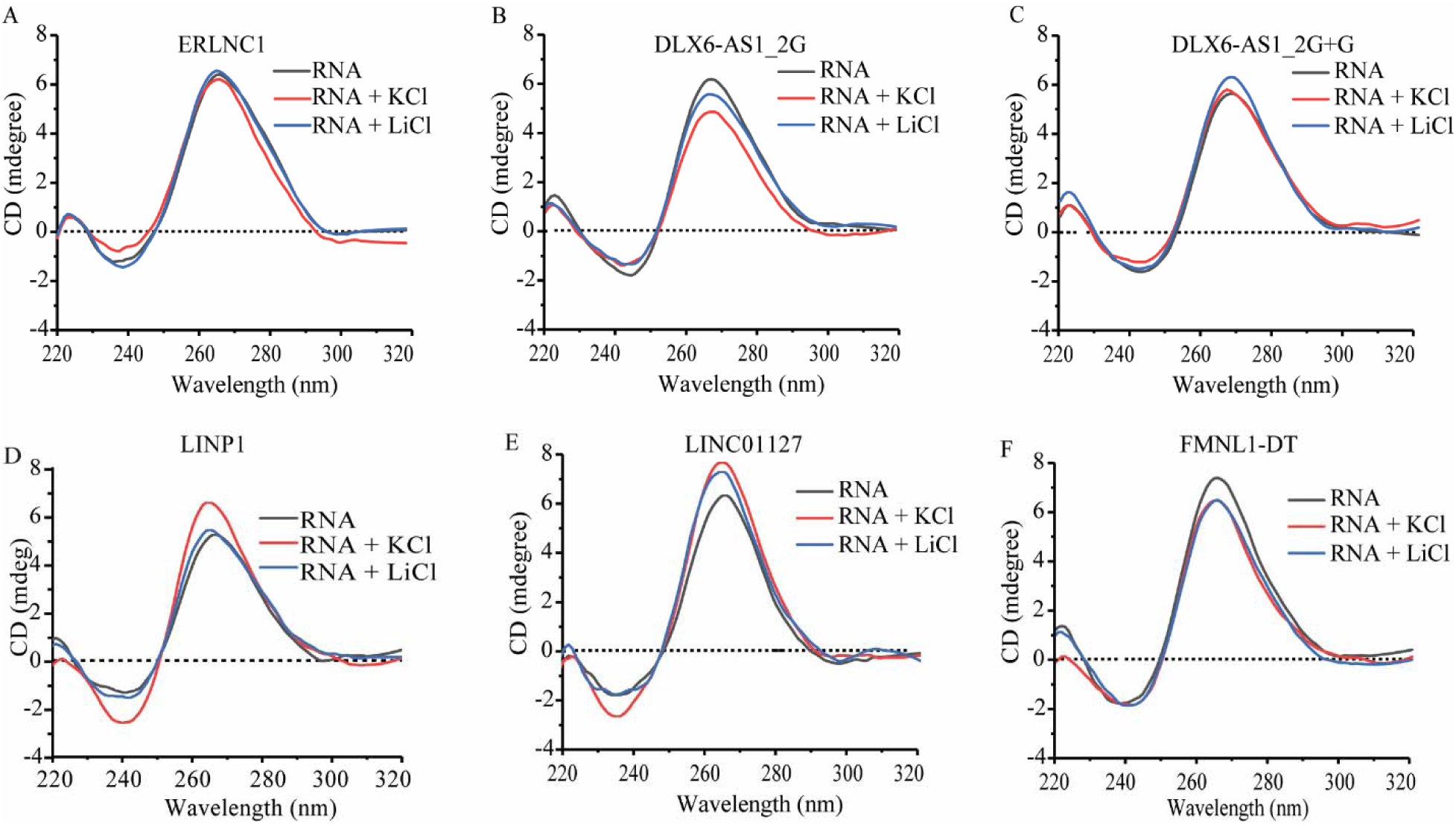
CD spectra of selected lncRNAs that are dysregulated in ovarian cancer. CD spectra of lncRNAs (5 µM) in the absence of additional ions and presence of monovalent cations K^+^ (100 mM) or Li^+^ (100 mM) (A) ERLNC1, (B) DLX6-AS1_2G, (C) DLX6-AS1_2G+G, (D) LINP1 (E) LINC01127 (F) FMNL1-DT.

**Fig. 2.**
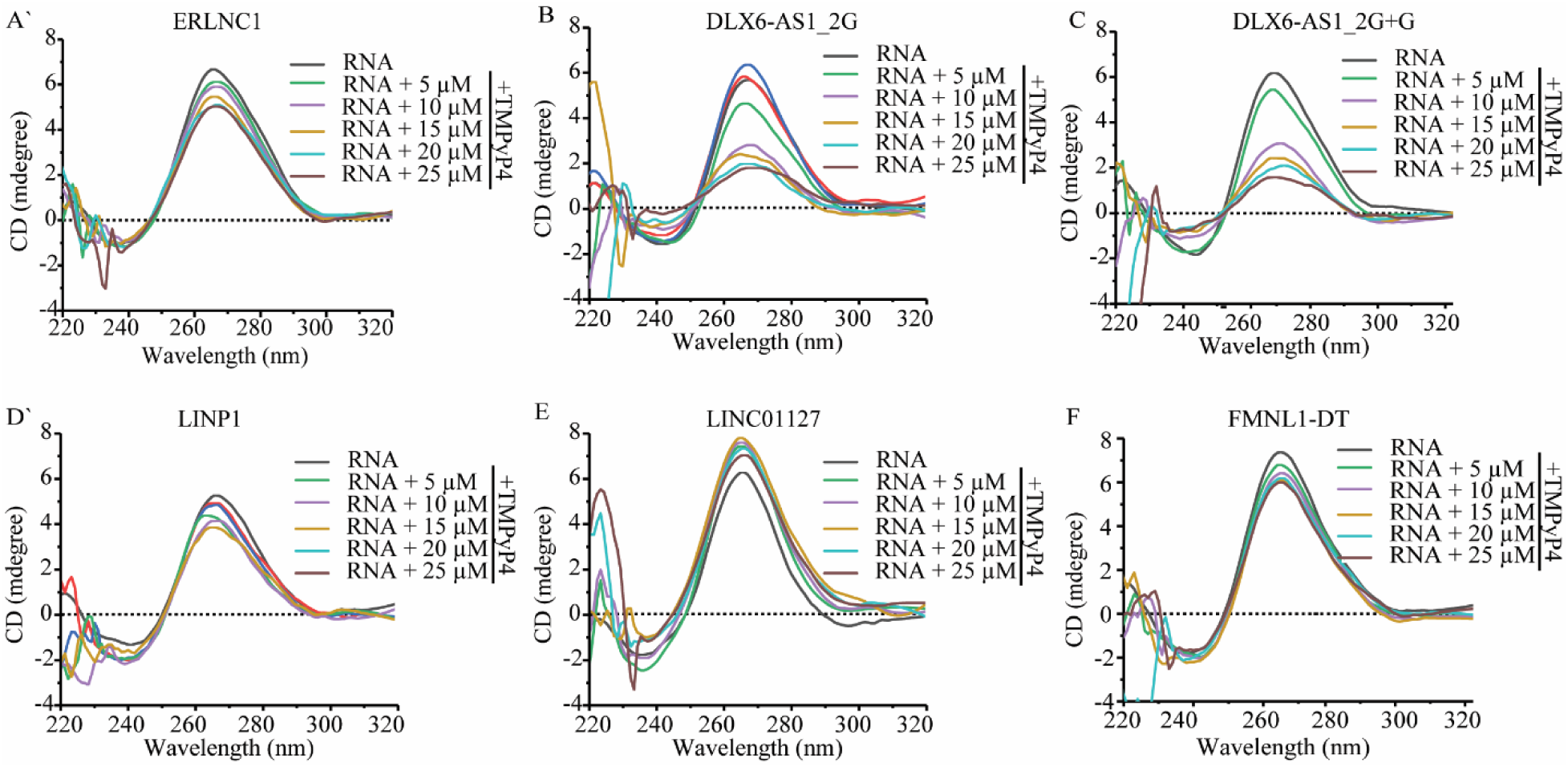
CD spectra of lncRNA G4s in varying amounts of TMPyP4. CD of lncRNAs (5 µM) in the presence of 5, 10, 15, 20, and 25 µM of TMPyP4. (A) ERLNC1, (B) DLX6-AS1_2G, (C) DLX6-AS1_2G+G, (D) LINP1 (E) LINC01127 (F) FMNL1-DT.

### 3.3. ThT fluorescence enhancement by RNA G4

Thioflavin T (ThT) fluorescence enhancement has been used to characterize RNA G4s(*39*). ThT demonstrates selective and preferential affinity for RNA G4s, resulting in enhanced fluorescence intensity when it binds to RNA G-quadruplex structures. A 50, 70, 140 and 200-fold of ThT fluorescence enhancement was obtained for the ERLNC1, DLX6-AS1_2G, DLX6-AS1_2G+G, and LINC01127 lncRNAs, respectively, with supplementation of KCl. In the presence of K^+^, FMNL1-DT and LINP1 displayed the greatest enhancement of ThT fluorescence of ∼260-fold and ∼270-fold, respectively (**Fig. 3G**). The fold of ThT fluorescence enhancement in presence of these RNAs can be contextualized by the nearly 90-fold enhancement in fluorescence observed for the lncRNA TERRA that was previously recorded by our lab(*21*). The concentration of the RNAs used for the ThT fluorescence experiments was two and a half-fold lower compared to that used for CD experiments described in the previous section. This was primarily to address the greater sensitivity of the ThT fluorescence assay compared to CD spectral measurements. The fold of ThT fluorescence enhancement for all lncRNAs, were significantly lower in presence of Li^+^. These results indicate that K^+^ consistently exerted a G4-stabilizing effect on the lncRNAs under study, albeit to a variable extent in each case **(Fig 3G)**. Notably, lncRNAs having higher fold enhancements, i.e., LINP1, LINC01127 and FMNL1-DT, are 3G-G4 forming sequences. 2G-G4 forming lncRNAs have comparatively lower fold enhancement as compared to 3G-G4s suggesting higher stabilities of the 3G-G4s, as intuitively expected.

**Fig. 3.**
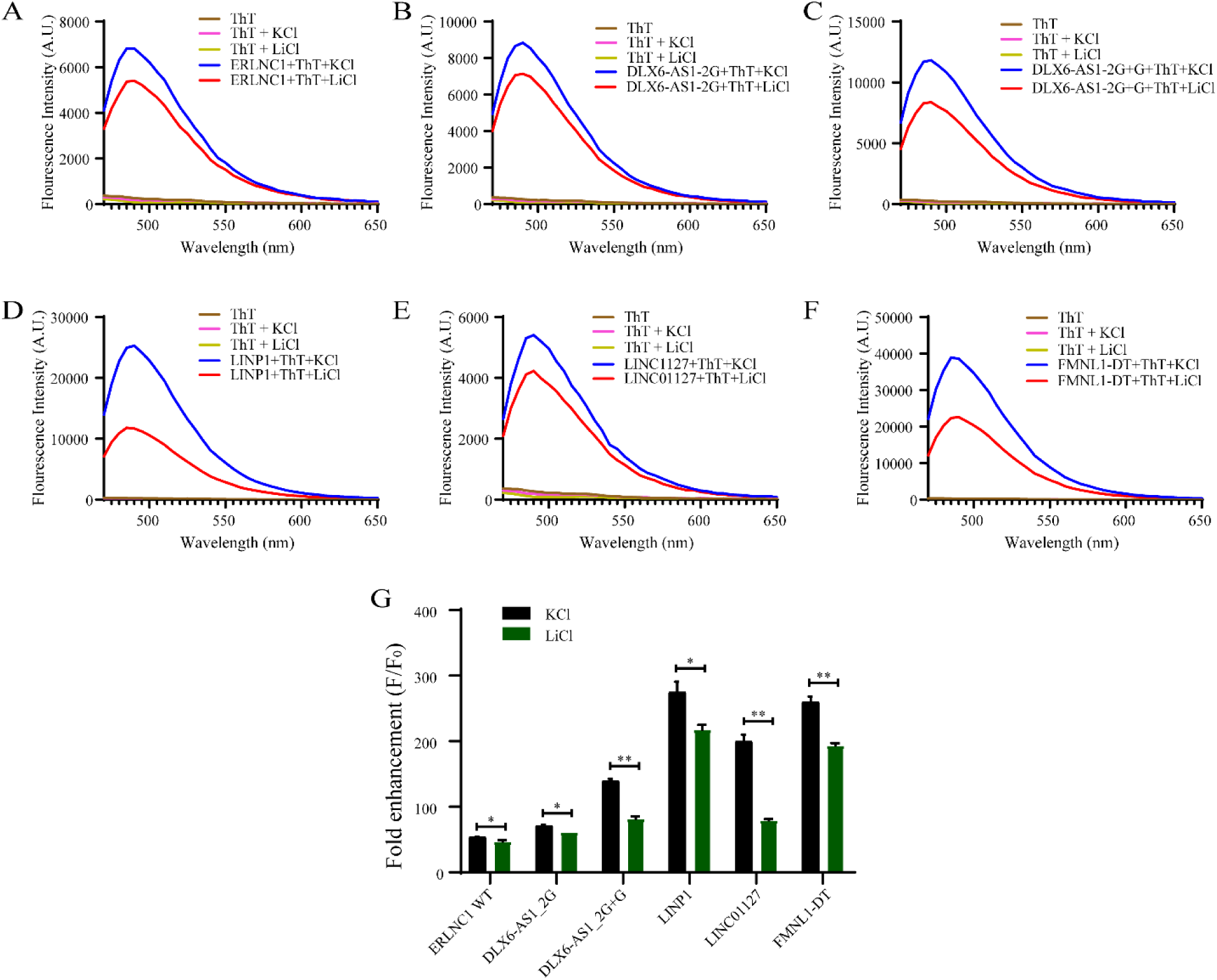
ThT fluorescence enhancement assay. Emission spectra of lncRNA (2 μM) in the presence of K^+^ or Li^+^ and ThT (2 µM) when excited at 445 nm. (A) ERLNC1, (B) DLX6-AS1_2G, (C) DLX6-AS1_2G+G, (D) LINP1 (E) LINC01127 (F) FMNL1-DT. (G) Fold enhancement of ThT fluorescence in the presence monovalent cations, with excitation and emission at 445 nm and 488 nm, respectively. Unpaired t-test was used for statistical analysis and the resulting statistical significance (*P*-values) are denoted with asterisks (*). Non-significant *P*-values are not represented.

We also examined the contribution of each G-tract towards maintaining the structural integrity of the corresponding G4s within each lncRNA. G-tracts were deleted in sequential manner to generate a set of deletion mutants. These mutants are designated as Δ1 (with the first G-tract removed), Δ2 (with second G-tract removed), and so on. Accordingly, we prepared six deletion mutants for ERLNC1, five for LINP1 and LINC01127, and four for DLX6-AS1. We were unable to investigate the effect of deletion of G-tracts on FMNL1-DT and ERLNC1_Δ1 due to challenges with *in vitro* transcription of the corresponding sequences. The IVT-derived wild-type (WT) lncRNAs and deletion mutants (Δ), lacking individual G-tracts, were analysed on a 15% native PAGE gel stained with ThT **(Suppl. Fig. 3).** For each lncRNA, the WT bands displayed the highest intensity, while the deletion mutants showed relatively weaker signals after ThT staining. These findings highlight the crucial role of specific G-tracts in maintaining the stability of the corresponding G4 structures. Scrutiny of the ThT fluorescence enhancement for the wild type versus deletion mutants across the different lncRNAs clearly indicates two features (**Fig. 4**). Firstly, the difference in fold-enhancement of ThT fluorescence between the WT and deletion mutants varies dramatically across the lncRNAs. Thus, the WT sequences of LINP1 and LINC01127 display significantly greater ThT fluorescence than the respective deletion mutants, and in the presence of both K^+^ and Li^+^. In contrast, the difference in fold enhancement between WT and deletion mutants of ERLNC1 and DLX6-AS1 is attenuated.

**Fig. 4.**
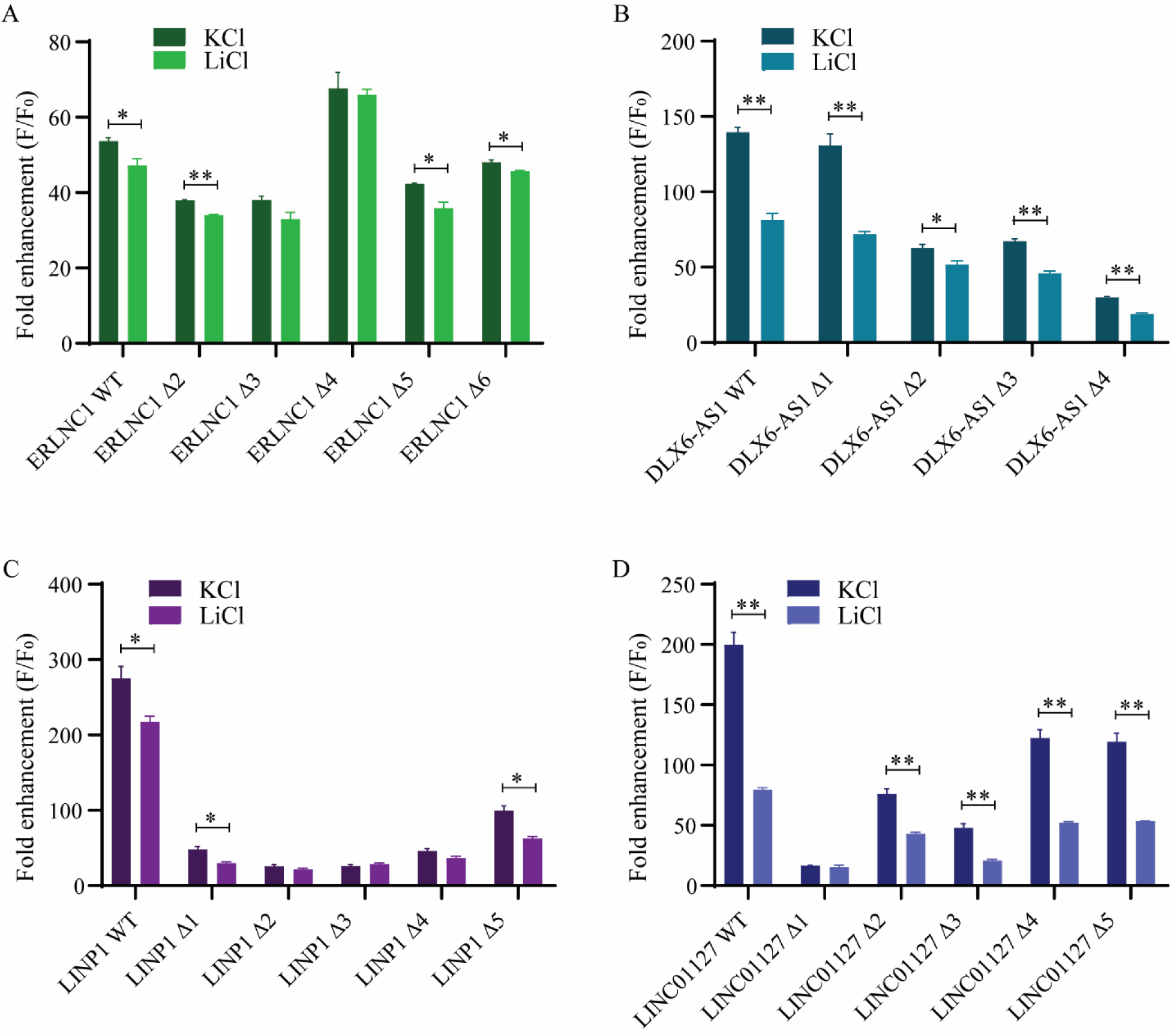
ThT fluorescence enhancement assay of deletion mutants. ThT fluorescence enhancement assay of wild-type RNAs and deletion mutants (2 µM) folded in the presence of 100 mM KCl or LiCl with ThT (2 µM). (A) ERLNC1, (B) DLX6-AS1_2G+G, (C) LINP1 (D) LINC01127. Fold enhancement of ThT fluorescence plotted in mean ± SEM at 488 nm when excited at 445 nm. Unpaired t-test was used for statistical analysis and the resulting statistical significance (*P*-values) are denoted with asterisks (*). Non-significant *P*-values are not represented.

Secondly, the removal of specific G-tracts results in the extremely sharp loss of fold of ThT fluorescence enhancement. For example, the removal of 2^nd^ G-tract of LINP1, 1^st^ G-tract of LINC01127 and 4^th^ G-tract of DLX6-AS1, result in greatest decrease in fold of ThT fluorescence enhancement compared to the cognate WT. Such reduction in the fold of ThT fluorescence can be correlated with the favourable contribution of the missing G-tract towards G4 formation by the WT sequence. Thus, the 2^nd^, 3^rd^ and 5^th^ G-tracts of ERLNC1, 2^nd^, 3^rd^ and 4^th^ G-tracts of DLX6-AS1, and all G-tracts for LINP1 and LINC01127 are essential for maintaining the stabilities of the respective G4s. While the absolute fold of ThT fluorescence enhancement for nearly all the WT and deletion mutants is greater in presence of K^+^ compared to Li^+^, the behavior of the deletion mutants vis-à-vis WT retains the same pattern between the two monovalent ions **(Fig. 4).**

### 3.4. Complementary DNA binding inhibits rG4 formation

We also assessed the contributions of individual G-tracts towards G4 folding by the respective lncRNA via a competitive binding assay of complementary DNA oligonucleotides. This assay relies on the hybridization of complementary DNA probes to a specific G-tract within the lncRNA, thereby blocking the constituent guanines from associating as G-quartets ultimately destabilizing the overall G4 structure. The designed competitive DNA oligos are complementary to individual G-tracts and the subsequent loop region. Incubation of the lncRNAs with DNA probes was followed by separation on native polyacrylamide gel and staining with ThT. This assay helps identify the G-tracts critical for G4 formation by observing the competitive inhibition of G4 formation as inferred from lowering of ThT fluorescence. We performed the complementary DNA binding assay on ERLNC1, DLX6-AS1_2G+G and LINP1. These sequences represent 2G and 3G sequences that display weak, modest and very strong effects of the respective deletion mutants. The results of the complementary DNA-mediated G4 inhibition assay are shown in **Fig. 5**. A close scrutiny of these results suggests that the effect of complementary DNA-binding could be placed under three categories: (1) increasing concentrations of the complementary DNA probe does not cause any change in G4 folding, (2) complementary DNA probe results in reduction in G4 folding but only when deployed at higher concentrations, and (3) complementary DNA results in significant loss of G4 folding across concentrations that are used. Accordingly, ERLNC1 is clearly a member of the first category, with some change in G4 formation observed for only one of the complementary DNA probes (ERLNC1:Comp4) at the highest concentration. Similarly, DLX6-AS1_2G+G belongs to the second category with respect to probes that are complementary to the 3^rd^ and 4^th^ G-tracts (DLX6-AS1:Comp3 and DLX6-AS1:Comp4). Interestingly, the fully complementary DNA probe (Comp1234) produces negligible changes in G4 folding by ERLNC1 in sharp contrast to the dramatic loss of G4-formation for DLX6-AS1_2G+G (**Fig. 5A-E, and 5F-J)**. For ERLNC1, these results could also be interpreted as the inability of the partially or fully complementary probes to bind with the G4 formed by the RNA. Notably, TMPyP4 was unable to exert meaningful disruption of the G4 formed by ERLNC1, supporting the robust structural features of the quadruplex (**Fig. 2**). In contrast, the PQS of DLX6-AS1_2G+G and LINP1 exhibited significant inhibition of G4 formation when the full-length complementary DNA oligo was applied, as indicated by a marked decrease in band intensity. LINP1 is mostly a member of the third category, with nearly all the probes resulting in reduction in G4 formation across all concentrations. The results of complementary DNA binding assay for ERLNC1, DLX6-AS1_2G+G and LINP1 are consistent with almost the entirety of results obtained for the deletion mutants of those RNAs. These findings underscore the crucial role of specific G-tracts within these lncRNAs in stabilizing their corresponding G4 structures.

**Fig. 5.**
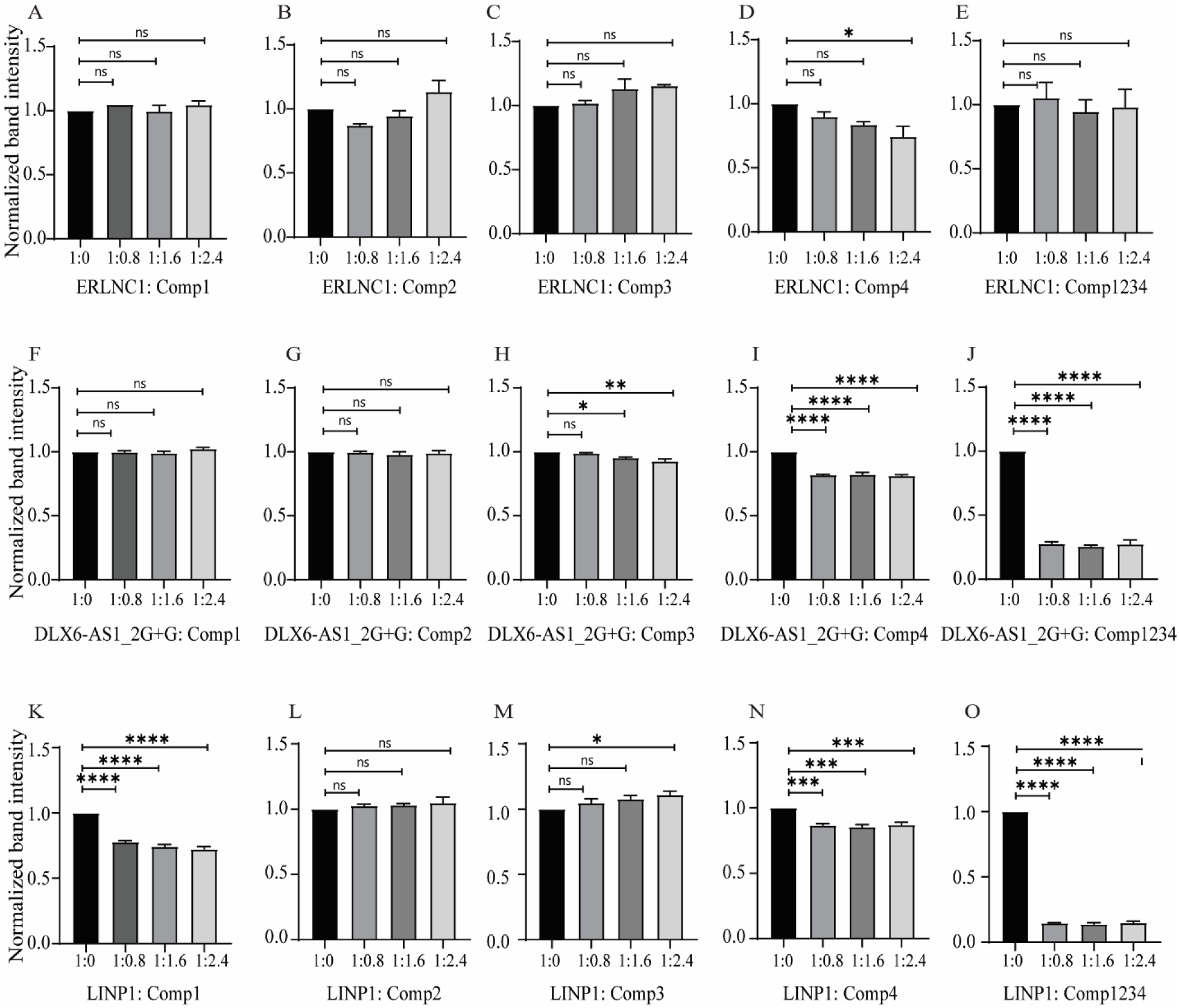
Competitor Oligonucleotide Mediated-G4 Disruption. Folded lncRNA PQS (2 µM) in the presence of increasing concentrations of DNA competitor, stained with 0.5 µM ThT. (A-E) ERLNC1 with Comp1, Comp2, Comp3, Comp4 and Comp1234 respectively. (F-J) DLX6-AS1_2G+G with Comp1, Comp2, Comp3, Comp4 and Comp1234 respectively. (K-O) LINP1 with Comp1, Comp2, Comp3, Comp4 and Comp1234 respectively. The bands were normalized against respective no-competitor control. *P*-values are calculated by ordinary one-way ANOVA and the resulting statistical significance are denoted with asterisks (*). Non-significant *P*-values are represented as ns.

### 3.5. Stabilization of RNA G4 structures by monovalent cations

We next used the Reverse Transcriptase (RT)-stop assay to investigate the response of G4s of ERLNC1, DLX6-AS1, LINP1, LINC01127 and FMNL1-DT to a processing enzyme, and shed light on their dynamic behaviour in presence of K^+^ versus Li^+^. The RT-stop assay is based on the interplay between the actively processing RT enzyme and the presence of G4 structures in the RNA template. The RT enzyme transcribes the RNA template until it reaches a stable RNA quadruplex structure. When quadruplex structures on the templates interfere with the RT process, truncated complement DNA products are produced and detected using denaturing PAGE assay(*40*). The effect of K^+^ and Li^+^ on the G4s can be assessed by inclusion of the monovalent ions in the RT-stop assay. The results of the RT-stop assay on ERLNC1, DLX6-AS1, LINP1 and FMNL1-DT are shown in **Fig. 6** and **Suppl. Fig. 4**. Two full-length products are observed in the RT-stop assay of ERLNC1, DLX6-AS1_2G, DLX6-AS1_2G+G, LINP1, FMNL1-DT and LINC01127, which could be due to the 5’ and 3’ heterogeneity in the RNA obtained via *in vitro* transcription(*41,42*). The results obtained from the RT-stop assay are discussed here vis-à-vis effects of K^+^ and Li^+^, and relative differences among the lncRNAs.

**Fig. 6.**
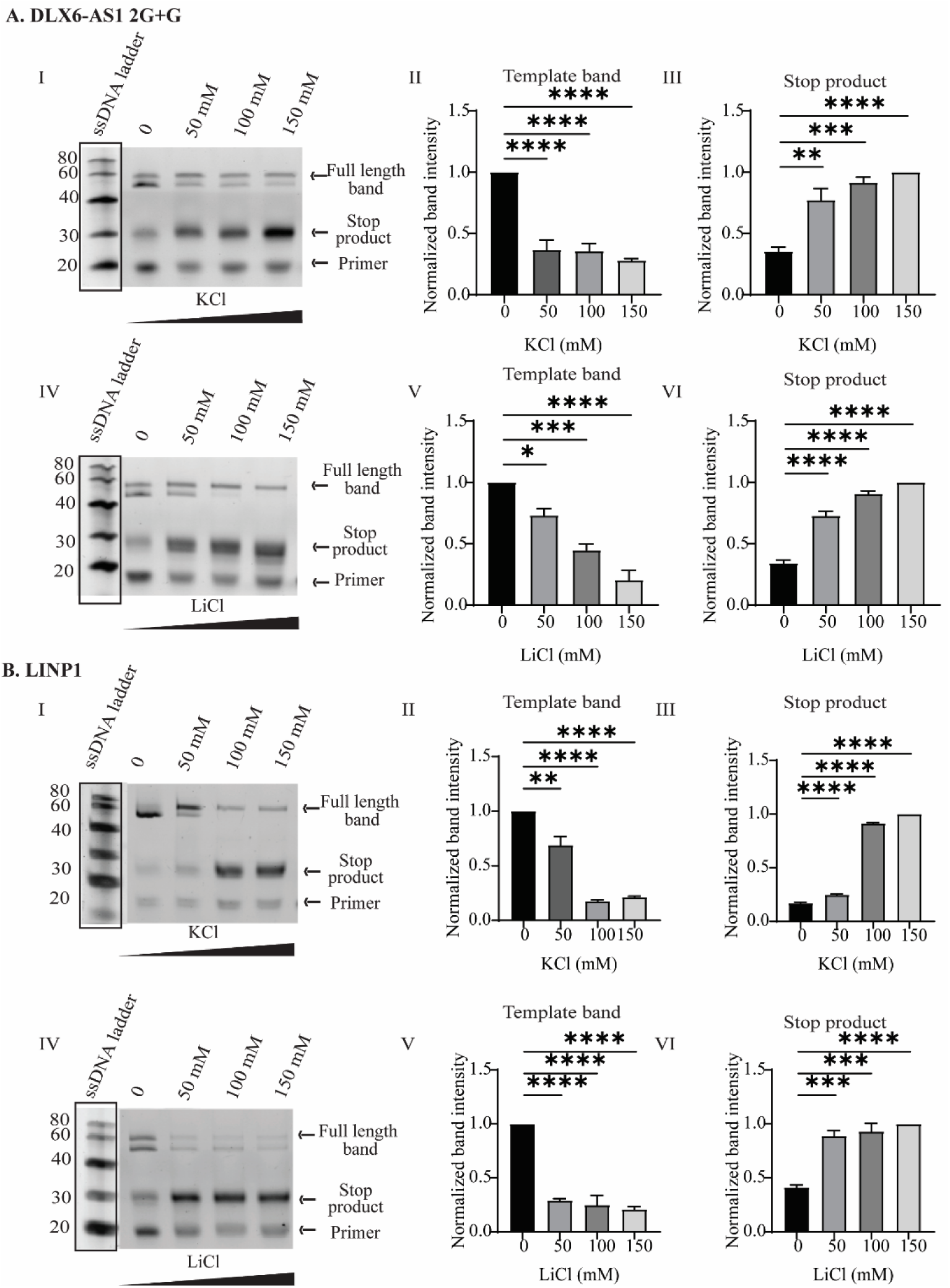
Reverse transcriptase stop assay for assessing influence of monovalent cations on the stability of G4s. A. DLX6-AS1_2G+G and B. LINP1 lncRNA in the presence of increasing concentrations of KCl and LiCl. **I.** shows the produced full-length (band 1 and 2) and truncated cDNA after reverse transcription of respective lncRNA in the presence of increasing concentrations (0, 50, 100, and 150 mM) of KCl *in vitro*. **II, III,** the correspondin quantification of full-length template bands and stop product **IV.** shows the produced full-length (band 1 and 2) an truncated cDNA after reverse transcription of lncRNA in the presence of increasing LiCl concentrations (0, 50, 100, and 150 mM) *in vitro*. **V, VI** is the corresponding quantification of full-length template bands and stop product. Ordinary one-way ANOVA was employed for the statistical analysis and the resulting statistical significance are denoted with asterisks (*). Non-significant *P*-values are represented as ns.

In the RT-stop assay of DLX6-AS1_2G+G, LINC01127 and FMNL1-DT, as the concentration of KCl increases, the production of truncated cDNA (stop products) increases, indicated by the corresponding stop products in the assay. This suggests that higher KCl concentrations stabilize the G4 structures of the respective lncRNAs, as these G4s cause reverse transcriptase to halt prematurely. The quantification of full-length bands (indicated by II in figure) reveals a gradual reduction in intensity, which correlates with an increase in truncated cDNA products (indicated by III in figure). Notably, different lncRNAs display varying degrees of sensitivity to KCl, with some lncRNAs (LINC01127) showing more prominent stops at lower KCl concentrations, while others (DLX6-AS1_2G+G FMNL1-DT) require higher concentrations to manifest significant stoppage of the RT. The RT-stop assay in the presence of LiCl for DLX6-AS1_2G+G, LINC01127 and FMNL1-DT depicts a different pattern of effects compared to KCl. Increasing concentrations of LiCl also lead to an increase in truncated cDNA, albeit in less pronounced manner compared to KCl. The quantification of full-length template bands (indicated by V in **Fig. 6**) and stop products (indicated by VI in **Fig. 6**) substantiates a similar trend, where the presence of LiCl stabilizes G4 formation but not to the same extent as KCl. The stop products increase in intensity, but the full-length bands persist longer compared to KCl assays.

This suggests that while LiCl stabilizes G4 structures, its effect is weaker than that of KCl for DLX6-AS1_2G+G, LINC01127 and FMNL1-DT. Interestingly, the lncRNAs LINP1, ERLNC1 and DLX6-AS1_2G exhibit a greater stabilizing effect from LiCl, with more distinct reverse transcription stops at lower concentrations of LiCl. The apparent stabilization of the G4s formed by LINP1, ERLNC1 and DLX6-AS1_2G in the presence of Li^+^ could be an indirect effect, arising from the suppression of alternative secondary structures. The capacity of cations to bind within the G-quartet stacks and influence G4 stability is influenced by the ion’s size, needing to be small enough to fit into the electron-rich cavity while also being large enough to connect the carbonyl oxygens of guanine(*43*). While K generally shows a greater stabilizing effect compared to Na and significantly greater than Li^+^, some exceptions exist for higher-order G4 structures(*44*). The effect of ion stabilization is a function of ion size and the precise guanine-tract architecture within each lncRNA, which dictates how well the G4 folds and stabilizes in the presence of specific ions(*45,46*). While LINP1, DLX6-AS1_2G and ERLNC1 appear to form G4s that are more attuned to Li ions, this may well reflect equilibria with non-G4 structures being more favored by K^+^ compared to Li^+^. The lncRNAs, DLX6-AS1_2G+G, LINC01127 and FMNL1-DT possess G-tract arrangements that engage favorably with K.

### 3.6. Dot Blot Assay validation of RG4 structures of OC-dysregulated lncRNAs

We were keen to examine the ability of the G4s formed by the OC-dysregulated lncRNAs to be recognized by G4-specific antibody BG4. The BG4 antibody selectively binds to various G-quadruplex types and does not bind to other nucleic acid motifs like RNA hairpins, single-stranded DNA, or double-stranded DNA(*47*). We used BG4 for dot blotting experiments with the *in vitro* transcribed lncRNAs containing PQS. The recognition of G4 by the antibody was assessed under different conditions such as monovalent ions, and G4-targeting ligands. The results of the dot blot assay are shown in **Fig. 7**. The RNA samples heated without EDTA were used as a “disabled sample” and RNA samples heated with EDTA were used as a “enabled sample.” The presence and absence of signals in enabled and disabled samples, respectively, reflect the facilitation or suppression of antigen behaviour of the nucleic acids. We used TERRA as a positive control in the dot blot assay based on its established RG4 motif(*21,48*). The results obtained by the dot blot assay offer a straight-forward interpretation of RG4 stability when exposed to K^+^/Li^+^ and different G4-targeting ligands. In presence of 100 mM KCl, ERLNC1, DLX6-AS1_2G+G, LINC01127, FMNL1-DT, and TERRA display strong BG4 signals, wherea LINP1 shows a moderate signal. In contrast, 100 mM LiCl results in weak BG4 signals across all lncRNAs except LINP1. This result is consistent with the higher stabilization of LINP1 by Li^+^ as observed in the RT stop assay.

**Fig. 7.**
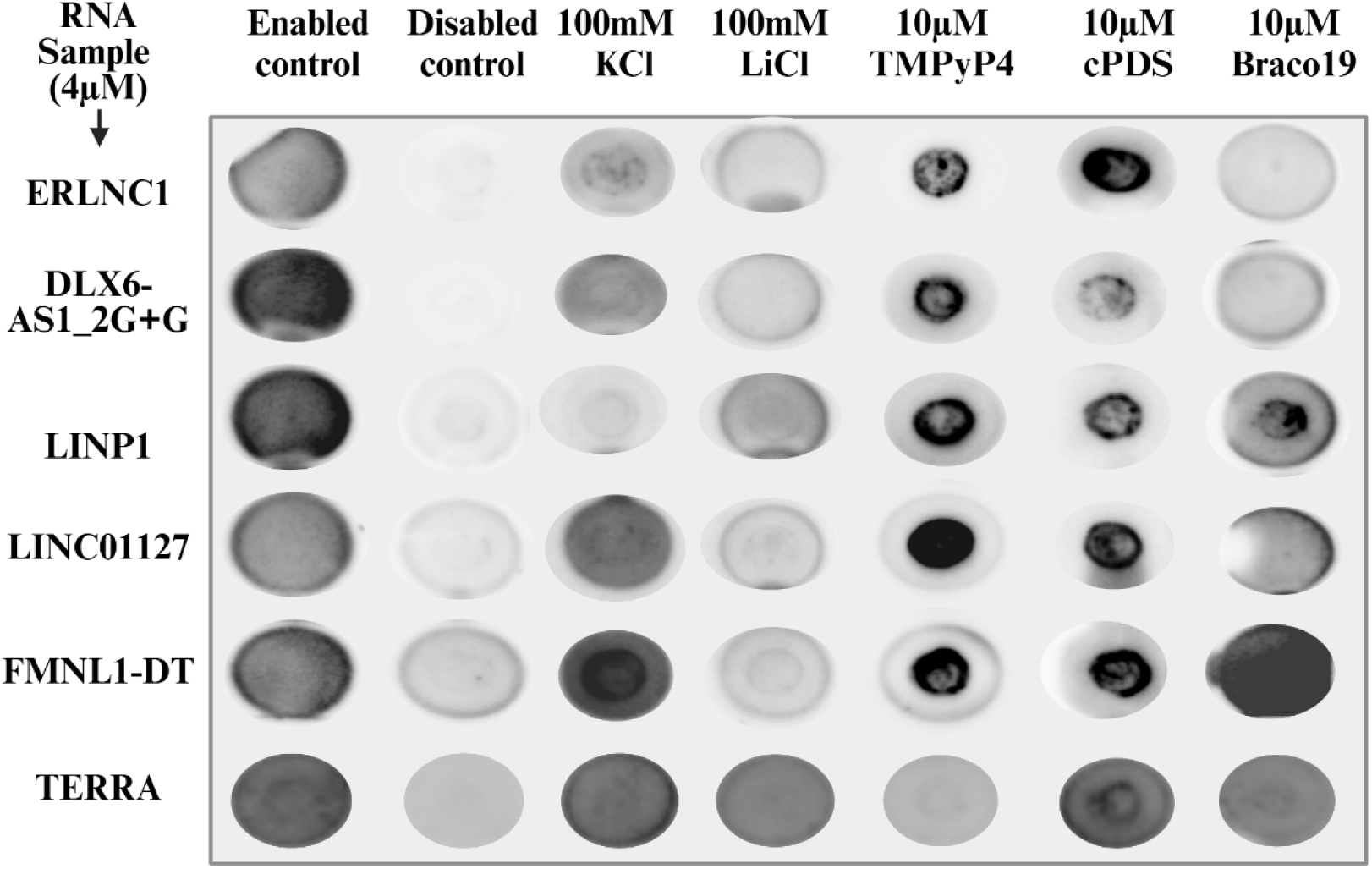
Dot blot assay for G4 forming lncRNAs in presence of different ions and ligands. Rows represent the lncRNAs ERLNC1, DLX6-AS1_2G+G, LINP1, LINC01127, FMNL1-DT, TERRA, and columns represent the treatment conditions: enabled control, disabled control, 100 mM KCl, 100 mM LiCl, 10 μM TMPyP4, 10 μM cPDS, and 10 μM Braco19.

In the presence of TMPyP4, a known RG4 destabilizer, a slight decrease in signal was observed. However, the concentration of TMPyP4 used (10 µM) was not sufficiently high to elicit significant destabilization of specific G4s as observed via CD spectroscopy (**Fig. 3**). Further, while all the lncRNAs showed positive signals in the presence of cPDS, Braco19 elicited weak signals from ERLNC1 and DLX6-AS1_2G+G. Notwithstanding the known RG4-stabilizing role of Braco19, the current results suggest subtle differences in the behaviour of the 2G versus 3G lncRNAs. Overall, the dot blot assay successfully demonstrates the formation and modulation of RG4 of various OC-dysregulated lncRNAs, as manifested by the binding of G4-specific antibody.

### 3.7. Human serum albumin interacts with G4 forming lncRNAs dysregulated in OC

In the previous section, we inferred the proteins interacting with RG4s of the OC-dysregulated lncRNAs studied in this work. As mentioned at the outset, we were keen to explore the behavioral facets of the RG4s in the context of OC-dysregulated circulating lncRNAs. In this regard, we were especially curious about the possible interactions between the RG4s and protein human serum albumin (HSA). HSA is the most abundant circulating protein found in plasma(*49*). HSA displays moderate binding affinity for various RNA molecules, including microRNAs such as miR4749 and miR-155. This interaction is primarily facilitated by electrostatic interactions between the negatively charged RNA and positively charged regions of HSA, along with hydrophobic forces from the hydrophobic regions of RNA(*50,51*). We investigated the interaction between HSA and the lncRNAs used in this study that are found in circulation, namely DLX6-AS1, LINP1, LINC01127. Our curiosity was based on the orthogonal roles reported for HSA with respect to RNA. Notably, HSA exhibits RNA-hydrolyzing activity, and possibly plays a role in RNA turnover within the bloodstream. While the RNA-hydrolyzing activity of HSA is significantly lower than that of ribonucleases, its potential involvement in the degradation of extracellular RNAs cannot be downplayed considering its high concentration in human blood(*52*). HSA has also been suggested as playing a protective role towards nucleic acids in the bloodstream and assist with the transport and delivery of nucleic acids to cellular targets(*53*). Interestingly, there is a dearth of reports on the interaction of circulating lncRNAs with HSA. We have used isothermal titration calorimetry (ITC), EMSA, and ThT fluorescence to investigate the interaction of HSA with ERLNC1, DLX6-AS1_2G+G and LINP1.

As presented in **Fig. 8**, ITC indicates thermodynamically favorable association between HSA and ERLNC1, DLX6-AS1_2G+G and LINP1. The ΔG for HSA association with ERLNC1, DLX6-AS1_2G+G and LINP1 were −9.42 kcal/mol, −9.87 kcal/mol, and −8.06 kcal/mol, respectively, indicating spontaneous binding. The K_D_ of HSA with ERLNC1, DLX6-AS1_2G+G, and LINP1 were estimated as 1.25 × 10^-7^ M, 5.82 × 10^-8^ M, and 1.25 × 10^-6^ M, respectively. The estimated K_D_ clearly points to high binding affinity for all the lncRNAs, especially for DLX6-AS1_2G+G. While ITC validates strong binding interactions between HSA and the RG4s, it is unable to unambiguously comment on changes experienced by the RG4 upon such binding.

**Fig. 8.**
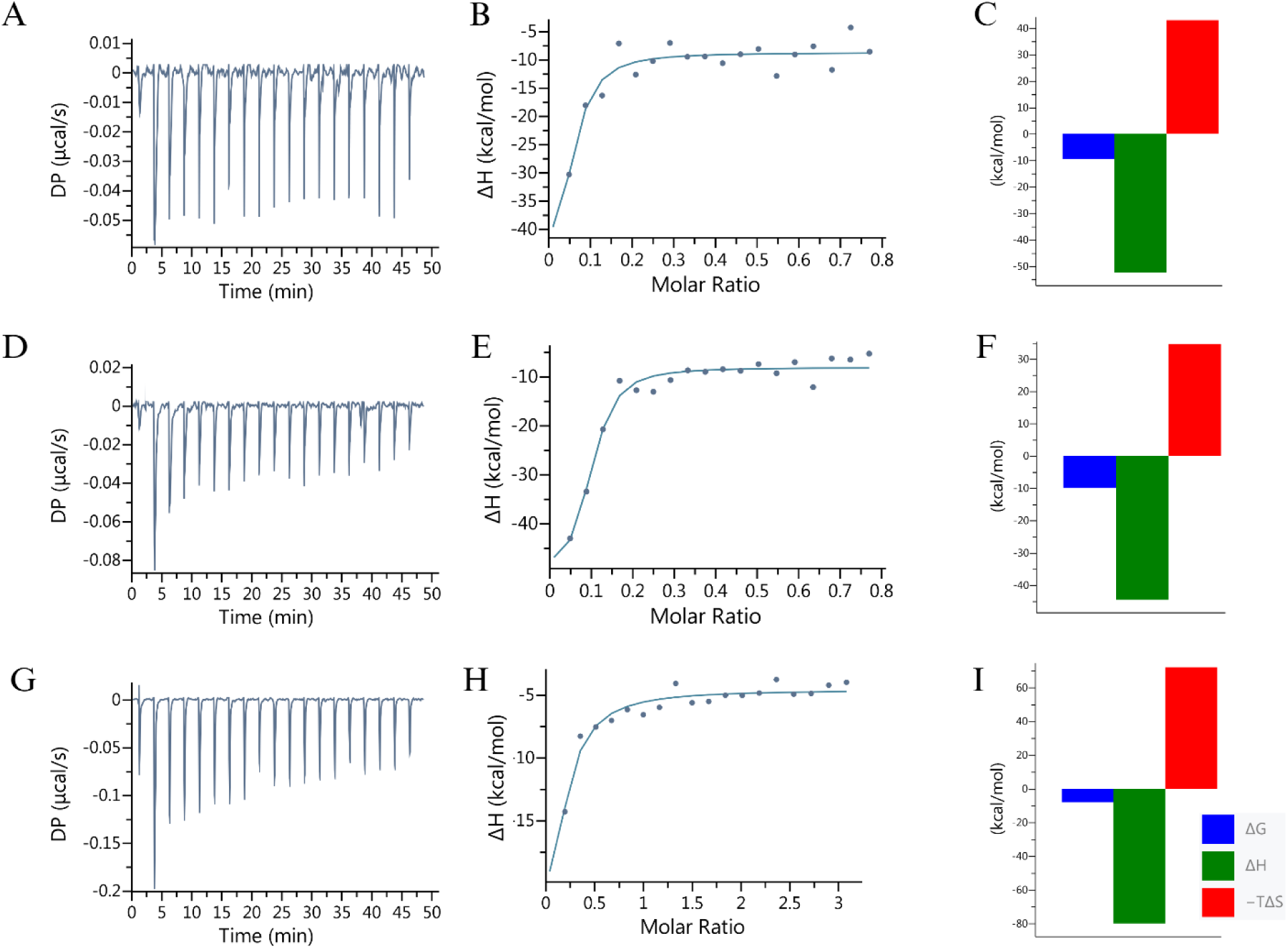
ITC of ERLNC1, DLX6-AS1_2G+G and LINP1 with HSA. Binding isotherms of, **A-C**. ERLNC1, **D-F.** DLX6-AS1_2G+G, **G-I.** LINP1, were fitted to determine ΔG, ΔH and TΔS for interaction of the RNAs with HSA.

Accordingly, we used ThT fluorescence enhancement to investigate the interaction of the RG4s with HSA, and specifically inform about the behavior of G4 motifs during the interaction. The ThT fluorescence enhancement assay was performed in increasing concentrations of HSA (5 - 60 µM). The fold of enhancement of ThT fluorescence decreased for all three lncRNAs with increasing amounts of HSA **(Fig. 9)**. This inverse relationship suggests that increasing concentrations of HSA could result in the following possible outcomes: (1) HSA displaces ThT from the G4-binding site based on its own affinity, and/or (2) HSA causes alteration in G4 conformation such that it no longer supports ThT binding. Both outcomes (1) and (2) would lead to lower fluorescence by weakening the basis of ThT fluorescence enhancement. Interestingly, the three lncRNAs display distinctly different patterns of change of ThT fluorescence. While increasing concentration of HSA into LINP1 results in a significant and progressive loss of ThT fluorescence, the addition of HSA to DLX6-AS1_2G+G results in loss of ThT fluorescence that is attenuated above 25 µM of HSA. The performance of ERLNC1 could be described as an in-between scenario, lying between the profiles of ThT fluorescence change observed for LINP1 and DLX6-AS1_2G+G. The significantly stronger K_D_ of DLX6-AS1_2G+G for HSA and the attenuated decrease in ThT fluorescence supports (1) formation of RG4-HSA complex at low concentrations of HSA, and (2) retention of G4 motif of the lncRNA even at high concentrations of the HSA. In other words, formation of a putative complex of DLX6-AS1_2G+G with HSA disturbs the G4 motif of the lncRNA to a limited extent. In contrast, the weaker K_D_ of LINP1 for HSA and the significantly greater loss of ThT fluorescence point to a severe disruption of the G4 motif of LINP1 upon association with HSA. ThT does not display meaningful fluorescence enhancement in presence of HSA alone thereby supporting the above-depicted behaviour of the lncRNAs.

**Fig. 9.**
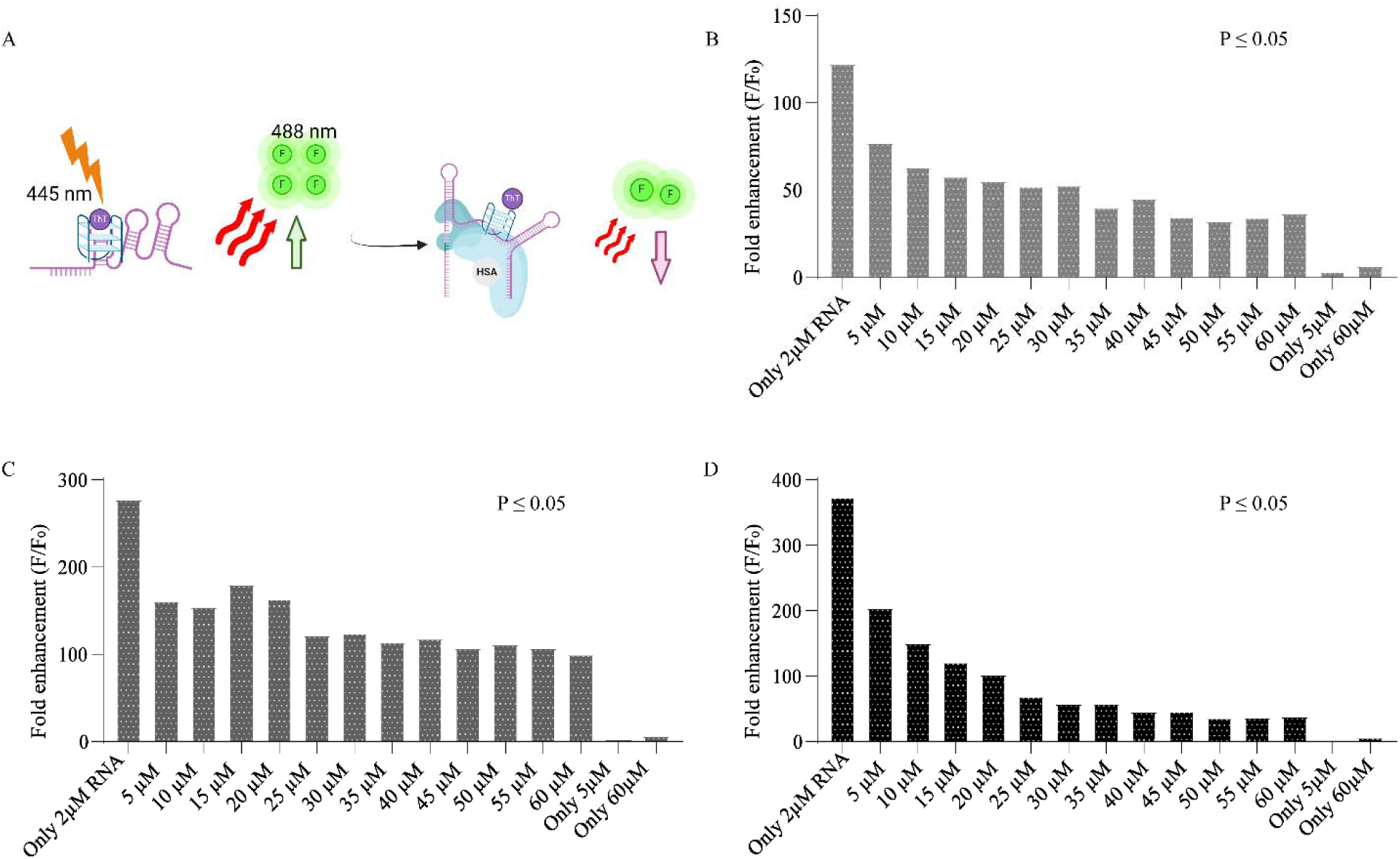
ThT fluorescence enhancement assay of ERLNC1, DLX6-AS1_2G+G and LINP1 in presence of HSA. (A) Schematic representation of the ThT fluorescence assay in the presence of lncRNAs and increasing concentration of HSA. (B-D) Fold enhancement of ThT fluorescence for rG4 lncRNAs (2 μM) (B) ERLNC1, (C) DLX6-AS1_2G+G, and (D) LINP1 in the presence of HSA (5 μM to 60 μM). Excitation and emission were at 44 nm and 488 nm, respectively. Unpaired t-test was used for statistical analysis and all the resulting *P*-values were significant.

The interaction of DLX6-AS1_2G+G with HSA points to possible changes in molecularity between the pristine RG4 and the RG4-HSA complex. We used EMSA to examine the potential changes in molecularity underlying the interaction of DLX6-AS1_2G+G with HSA. The lncRNA was labeled with Texas Red-labeled primer attached to the primer binding site. The primer-annealed RNA was incubated with different HSA concentrations and then loaded onto 8% polyacrylamide gels. The gels were imaged using the Rhodamine filter to visualize the presence of the primer-bound RNA. Counterstaining of the gels with ThT permitted visualization of the G4 forming species. In the presence of HSA, the relative mobility of the folded lncRNA (DLX6-AS1_2G+G) changed significantly **(Fig. 10A)**. Increase in HSA concentration resulted in formation of rapid moving bands. As shown in **Fig. 10A**, the intensity of band 1 decrease progressively while the intensity of the fast-moving product (band 2) increases with higher HSA concentrations. The ThT-stained gel is shown in **Fig. 10B** with the prospective G4-bearing species indicated as band 3. The quantification of bands 1, 2, and 3 is presented in **Fig. 10C, D, and E**, respectively. EMSA was conducted for other selected rG4s; however, all other sequence produced fast-moving products without evidence of G4 formation after ThT staining. The EMSA results for lncRNA LINP1 with HSA are shown in **Suppl. Fig. 5**.

**Fig. 10.**
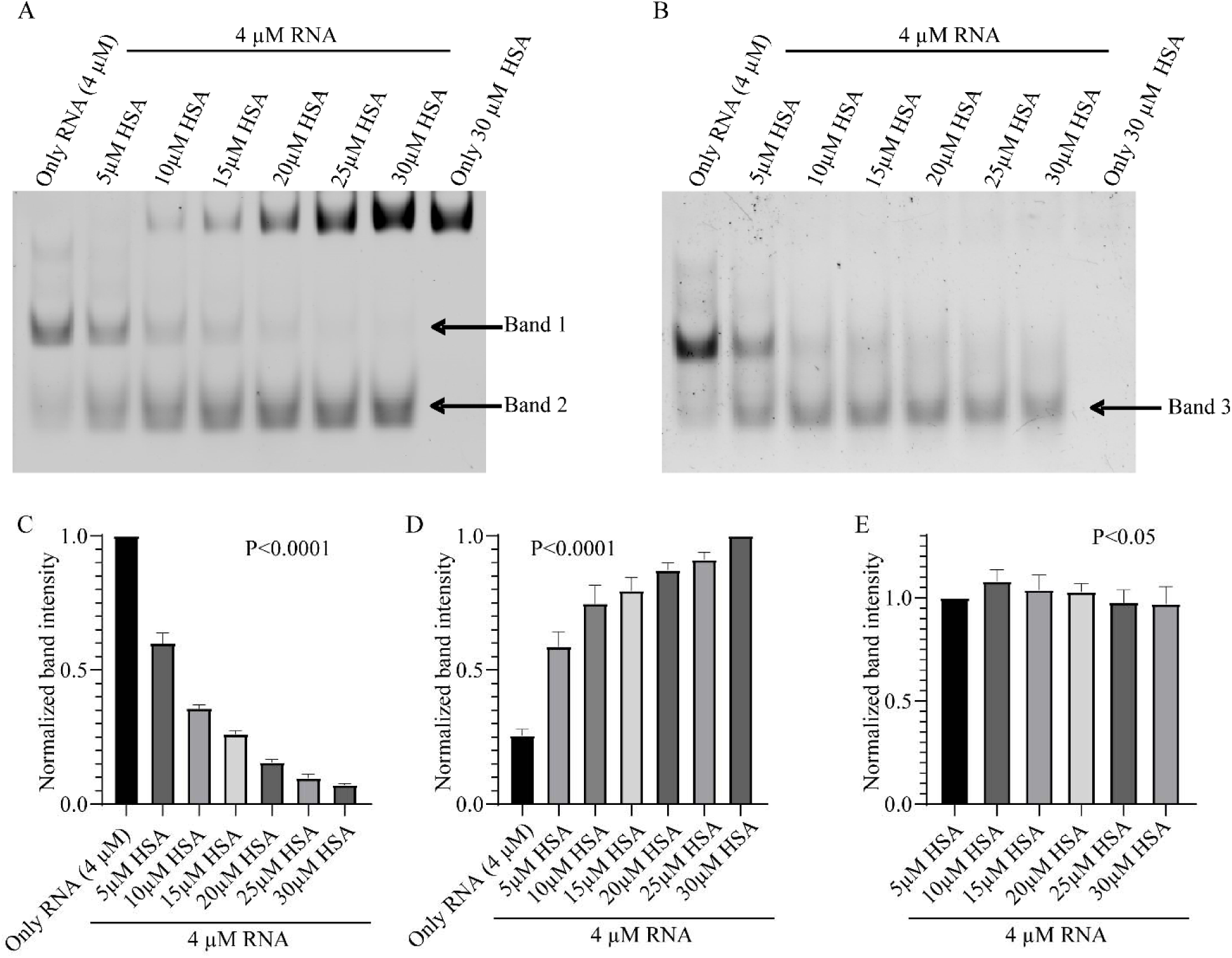
EMSA of lncRNA DLX6-AS1_2G+G with HSA. A. Annealed and folded DLX6-AS1_2G+G (4µM) with increasing concentration of HSA visualized in rhodamine filter B. Annealed and folded DLX6-AS1_2G+G (4µM) with increasing concentration of HSA stained with ThT. C, D, E. Quantification of Band 1, 2 and 3 respectively as indicated in figure. Ordinary one-way ANOVA was employed for the statistical analysis, and all the resulting *P*-values were significant.

The emergence of bands 2 (and 3) at low concentrations of HSA align with the strong binding affinity of the DLX6-AS1_2G+G for the protein. The EMSA of DLX6-AS1_2G+G with HSA points to: (1) formation of a rG4-protein complex that displays faster mobility compared to the native rG4, and (2) no significant disruption in the G4 motif of the lncRNA. These inferences are consistent with the previously described ITC and ThT fluorescence results. Nevertheless, the faster mobility of the lncRNA-protein complex is unusual and intriguing as the complex would have a higher molecular weight. The mobility of complexes during gel electrophoresis depends on several parameters, including molecular mass, charge, and conformation(*54*). It is possible that the slower migrating species belonging to the uncomplexed lncRNA corresponds to a multimeric intermolecular complex. Previous reports have attributed slower moving species to formation of multimeric intermolecular complexes (*55*). The binding affinity of HSA for DLX6-AS1_2G+G is sufficiently strong to disintegrate the multimeric complex of the RNA while enabling retention of the G4 motif in the resulting RNA-HSA complex. The remarkable interaction of DLX6-AS1_2G+G with HSA is worthy of further scrutiny to assess potential functional implications. Since some of the selected lncRNAs interact with HSA, we employed RNA-Protein Interaction Prediction (RPISeq) database, which identifies its most likely protein binding partner from a user-provided list for a given RNA sequence. RPISeq employs machine learning classifiers to generate interaction probabilities, ranging from 0 to 1. In performance evaluation, predictions with probabilities over 0.5 are rated as “positive,” indicating a strong potential for interaction between the corresponding RNA and protein. As indicated in **Table 3**, DLX6-AS1 shows highest binding potential with HSA, which is in corroboration with our experimental results.

**Table 3.**
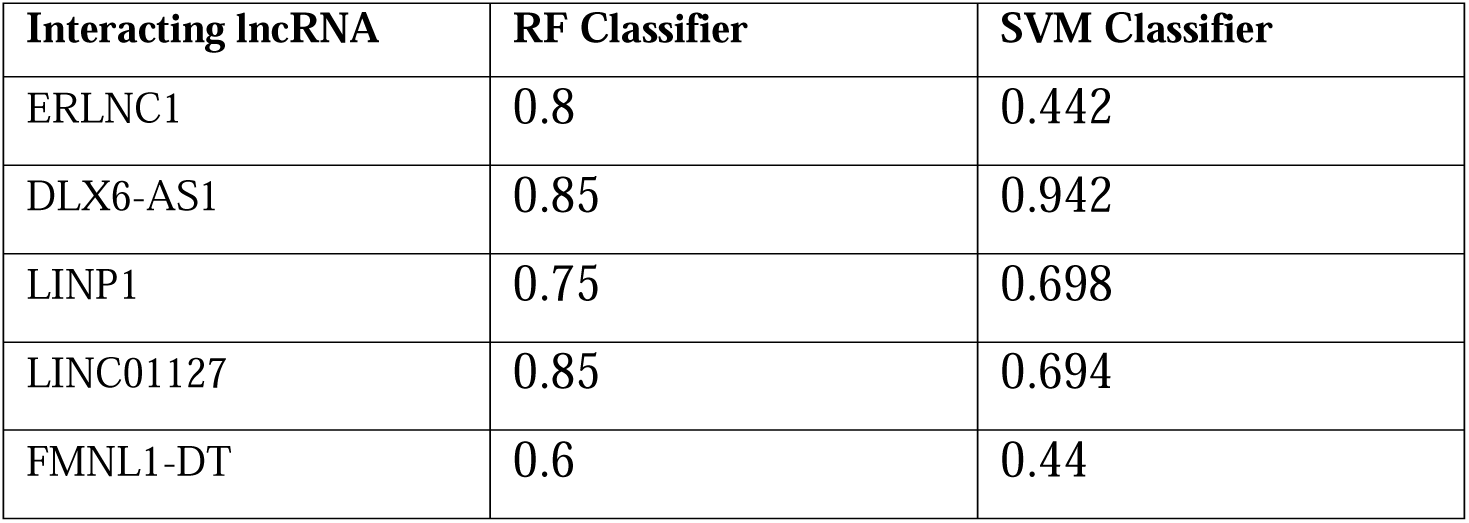
RPISeq predicted binding potential of lncRNAs to HSA protein.

| Interacting lncRNA | RF Classifier | SVM Classifier |
| --- | --- | --- |
| ERLNC1 | 0.8 | 0.442 |
| DLX6-AS1 | 0.85 | 0.942 |
| LINP1 | 0.75 | 0.698 |
| LINC01127 | 0.85 | 0.694 |
| FMNL1-DT | 0.6 | 0.44 |

## 4. Conclusion

The experimental results provide compelling evidence that dysregulated lncRNAs in ovarian cancer can form G-quadruplex (G4) structures, which play a crucial role in regulating gene expression and interaction with proteins. Using computational tools such as Lnc2Cancer, QGRS Mapper, and G4Hunter, four lncRNAs were identified with a high potential for G4 formation. These lncRNAs include DLX6-AS1, LINC01127, FMNL1-DT, and ERLNC1, with specific sequences exhibiting distinct G4 topologies confirmed by circular dichroism (CD) spectroscopy. The presence of parallel G4 structures was observed across all selected lncRNAs, with varying stability influenced by the presence of monovalent cations like KCl and LiCl.

The study also demonstrated that TMPyP4, a porphyrin-based ligand, effectively destabilizes the rG4 structures, leading to a conformational change, as evidenced by decreased CD spectra intensity. This suggests that targeting G4 structures with specific ligands could be a potential therapeutic strategy for modulating the function of these lncRNAs in ovarian cancer. Thioflavin T (ThT) fluorescence enhancement experiments demonstrate that K^+^ stabilizes rG4 structures in ovarian cancer-associated lncRNAs, with 3G-G4 forming sequences like LINP1, LINC01127, and FMNL1-DT showing the greatest fluorescence enhancement. Deletion mutant analyses reveal that specific G-tracts are essential for maintaining the structural integrity of G4s. Complementary DNA binding assays further support the critical role of these G-tracts in G4 formation, particularly in DLX6-AS1_2G+G and LINP1. These results highlight the importance of G-tract architecture in G4 stability and function. The RT-stop assay reveals that K^+^ significantly stabilizes rG4 structures in ovarian cancer-associated lncRNAs, particularly DLX6-AS1_2G+G, LINC01127, and FMNL1-DT, leading to increased reverse transcriptase halting. While Li^+^ also stabilizes G4s, its effect is generally weaker than K+, except in LINP1, ERLNC1, and DLX6-AS1_2G, where Li^+^ exerts a stronger influence. This differential stabilization may be due to the unique guanine-tract architecture within each lncRNA, which affects how well G4s fold and interact with specific cations. These findings underscore the role of cations in modulating G4 structure and stability. The dot blot assay confirms the formation of G4s, as detected by the G4-specific BG4 antibody. The stability of these G4s varies based on monovalent ions and G4-targeting ligands, with KCl generally enhancing G4 recognition, while LiCl shows weaker effects except for LINP1. The interaction studies between human serum albumin (HSA) and rG4s formed by OC-dysregulated lncRNAs reveal significant binding affinities, particularly with DLX6-AS1_2G+G. This interaction preserves the G4 structure, as evidenced by ITC, EMSA, and ThT fluorescence assays. While HSA binding impacts the molecularity of DLX6-AS1_2G+G, it does not significantly disrupt the G4 motif, indicating a stable complex. The findings suggest a functional role for HSA in stabilizing circulating lncRNAs, potentially influencing their stability and transport in the bloodstream. These insights warrant further exploration into the biological implications of HSA-RG4 interactions. Overall, this study underscores the importance of G4 structures in lncRNAs dysregulated in ovarian cancer, providing a foundation for future research into their biological roles and therapeutic potential.

## CRediT authorship contribution statement

Deepshikha Singh: Conceptualization, Investigation, Methodology, Data curation, Validation, Formal Analysis, Visualization, Writing – original draft, Writing – review & editing. Chinmayee Shukla: Conceptualization, Investigation, Methodology, Writing – review & editing. Bhaskar Datta: Conceptualization, Methodology, Validation, Writing – original draft, Writing – review & editing, Funding acquisition, Project administration, Resources, Supervision.

## Declaration of competing interest

None. The authors declare that they have no known competing financial interests or personal relationships that could have appeared to influence the work reported in this paper. Signed by corresponding author on behalf of all authors.

## Acknowledgements

B.D. gratefully acknowledges financial support for this work by Gujarat State Biotechnology Mission (GSBTM) vide project no. GSBTM/JD(R&D)/626/22-23/00006262. All the illustrations are created with Biorender.com.

## Notes

### Competing Interest Statement

The authors have declared no competing interest.

