## Supplementary Information for "G-quadruplex Structures in Dysregulated Long Non-Coding RNA of Ovarian Cancer and their Binding Interactions with Human Serum Albumin"

* Corresponding author.

**1. Oligonucleotides for *in vitro* transcription**

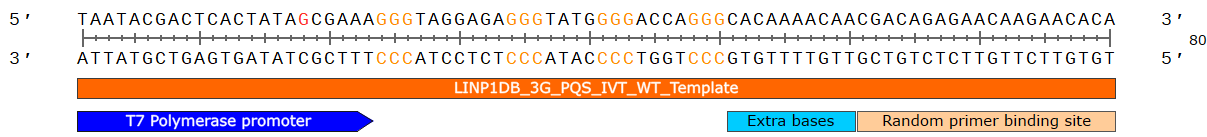

**Suppl. Fig. 1.** Oligonucleotide design for IVT

The cDNA antisense template was designed with SnapGene software (Dotmatics, [www.snapgene.com](http://www.snapgene.com/), USA). Design of DNA oligonucleotide template (bottom strand), consisting of antisense DNA strand of the T7 RNA promoter, the antisense of the PQS region of the respecftive lncRNA, followed by a few extra bases, and then the random primer binding site required for reverse transcription.

**2. T7 promoter Modification**

Considering the promoter sequence displayed below, *in vitro* transcription begins that the highlighted G. The polymerize transcribes from 5’ → 3’ using the opposite strand as a template. The presence of a G as the first base in the transcript could pose challenges for the investigation of G4 formation by the RNA. Notably, the PQS region of the lncRNA may include the first G to generate a G4. We have modified the T7 promoter sequence slightly to prevent the initial G from being added and resulting in disruption.

**Actual T7 Promoter sequence:**	5′ TAATACGACTCACTATAG 3′

**High yield T7 Promoter:	5’** TAATACGACTCACTATAGGG 3’

**Modified T7 promotor sequence:**	5’ TAATACGACTCACTATAGCGAAA 3’

Although it is desirable that the final template be created so that transcription begins with at least two Gs, there would be no way to identify which G tract is involved in G4 development since the two Gs would interfere with G4 formation and behave like another G tract. Low 5' heterogeneity and high yield are achieved by modifying the T7 promoter sequence.

**3. Experiment design of competitive binding of complementary DNA oligos**

The competition assay is based on the competitive binding of a complementary DNA oligonucleotide to a specific G-tract of the PQS of the *in vitro* synthesized lncRNA sequence. Competitive DNA oligos are designed in such a way that they are complementary to a specific G-tract and the subsequent loop region. This assay sheds light on the G-tracts that contribute significantly towards G4 formation, based on the competitive inhibition due to hybridization of the complementary DNA sequences.

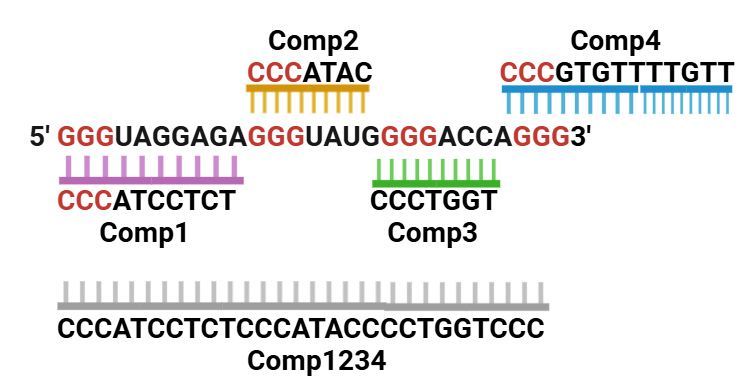

**Suppl. Fig. 2.** Design of DNA oligonucleotides for the competition assay.

**4. Native PAGE of wild type and deletion mutants of selected lncRNAs**

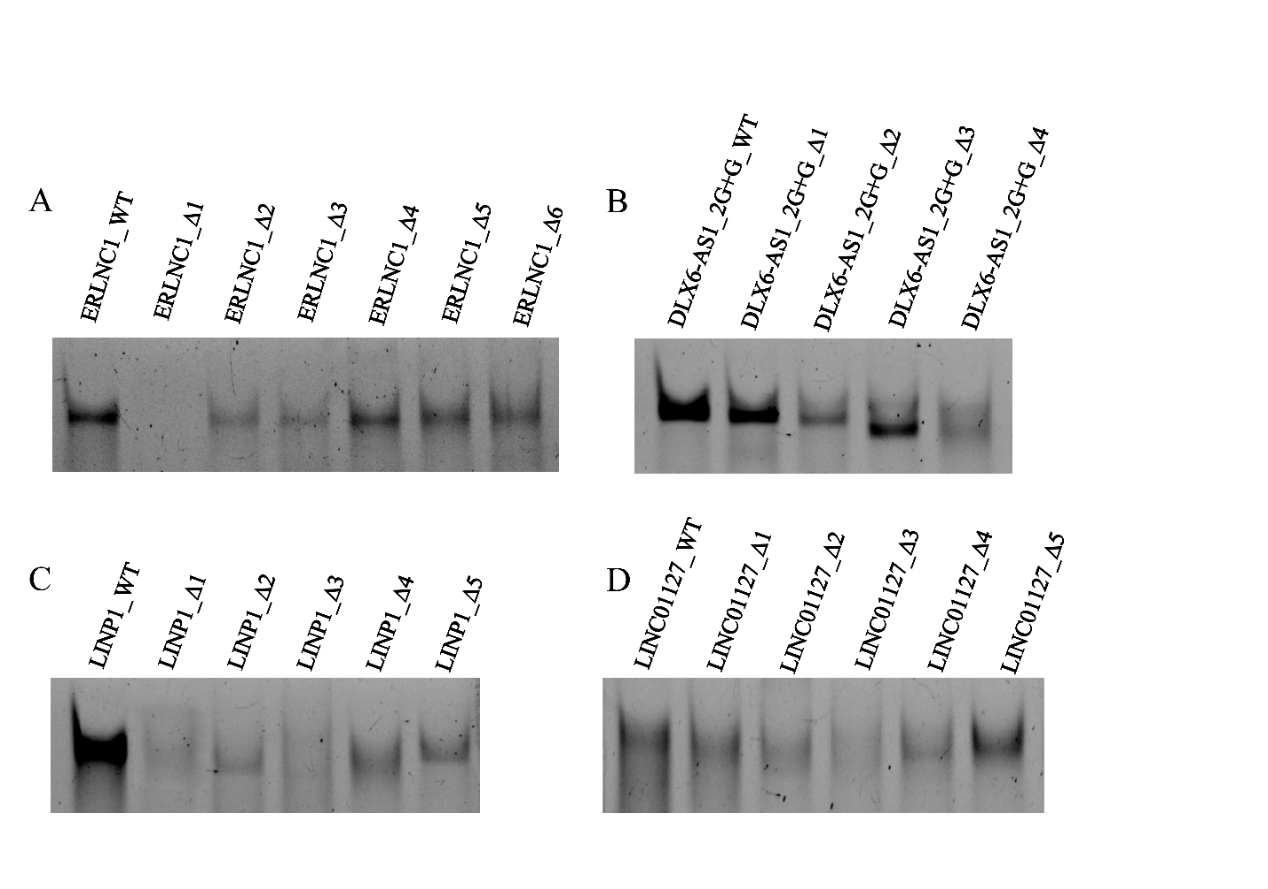

**Suppl. Fig. 3. G-tracts influence the stability of lncRNA G4s.** Native PAGE (15%) of folded IVT-derived wild-type and deletion mutants of RNAs (2 µM) stained with ThT (0.5 µM). A. ERLNC1 B. DLX6-AS1_2G+G C. LINP1 D. LINC01127.

**5. Reverse transcriptase stop assay of selected lncRNAs**

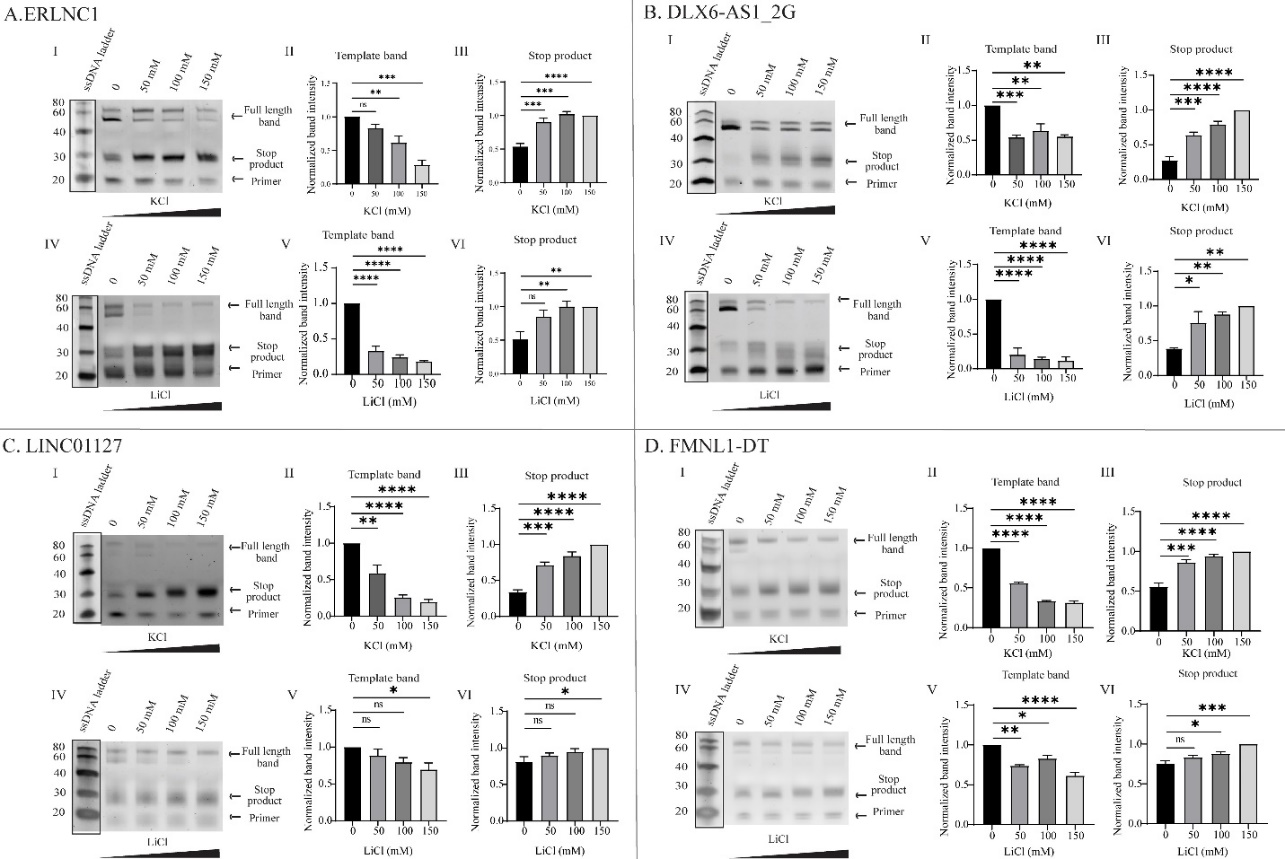

**Suppl. Fig. 4.** Reverse transcriptase stop assay of A. ERLNC1 B. DLX6-AS1_2G C. LINC01127 D. FMNL1-DT lncRNA in the presence of increasing concentrations of KCl and LiCl. **I.** shows the produced full-length (band 1 and 2) and truncated cDNA after reverse transcription of respective lncRNA in the presence of increasing concentration (0, 50, 100 and 150 mM) of KCl *in vitro*. **II, III** the corresponding quantification of full-length template bands and stop product **IV.** shows the produced full-length (band 1 and 2) and truncated cDNA after reverse transcription of lncRNA in the presence of increasing concentration (0, 50, 100, and 150 mM) of LiCl *in vitro*. **V, VI** the corresponding quantification of full-length template bands and stop product. Ordinary one-way ANOVA was employed for the statistical analysis and the resulting statistical significance are denoted with asterisks (*). Non-significant *P*-values are represented as ns.

**6. EMSA of lncRNA LINP1 with HSA**

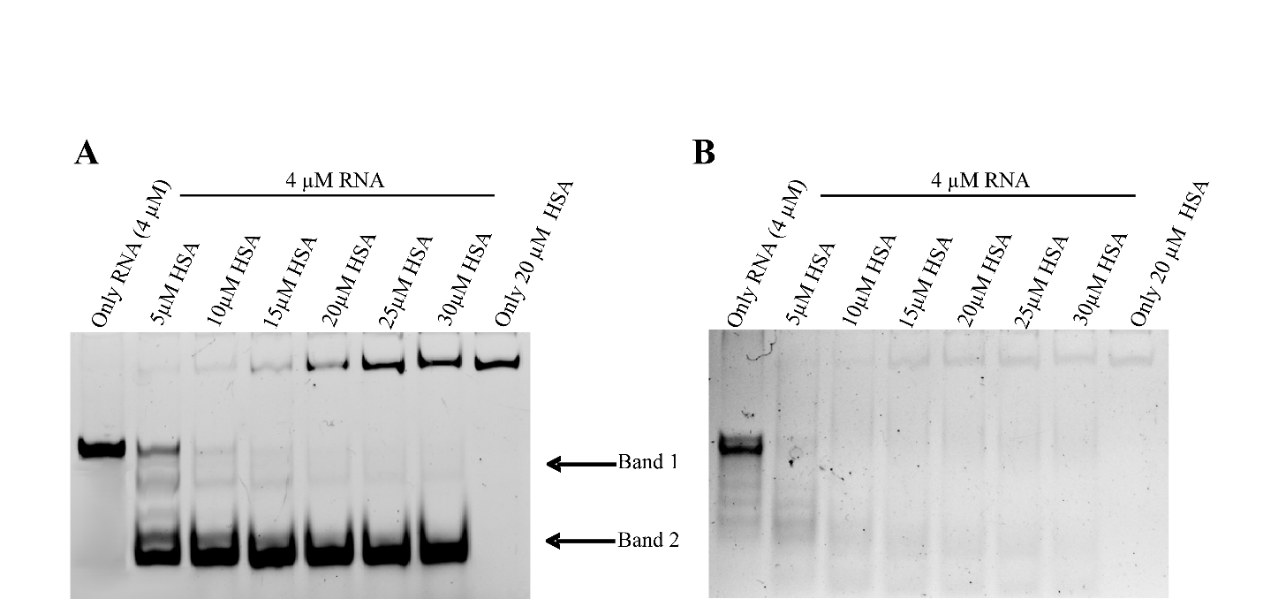

**Suppl. Fig. 5: EMSA of lncRNA LINP1 with HSA.** A. Annealed and folded LINP1 (4µM) with increasing concentration of HSA visualized in rhodamine filter B. Annealed and folded LINP1 (4µM) with increasing concentration of HSA stained with ThT.
